# RNA polymerase loss by nuclear rupture drives *LMNA* cardiomyopathy

**DOI:** 10.64898/2026.04.03.716433

**Authors:** Atsuki En, Melanie Gucwa, Erjola Rapushi, Cara Barnett, Wataru Katano, Nonso Nduka, Tatsuya Shiraki, Alyssa Grogan, Aloke V. Finn, K. Nicole Weaver, Kohta Ikegami

## Abstract

Localized rupture of the nuclear envelope has recently been reported in various pathologies, including cancer ^1,2^, neurodegenerative disease ^3–5^, myocardial infarction ^6^, as well as dilated cardiomyopathy caused by Lamin A/C gene mutations (LMNA-DCM) ^7^. Whether and how nuclear rupture contributes to disease remains unknown. Here, we report that nuclear rupture causes global transcriptional deficiency in a mouse model of LMNA-DCM. We observed that ruptured nuclei lost RNA polymerase II, leading to downregulation of numerous genes essential for cardiomyocyte structure and function. We identified endogenous resealing of nuclear rupture as a cardioprotective mechanism in LMNA-DCM mouse hearts. Resealing involved the ESCRT-III membrane remodeling complex recruited to nuclear rupture sites. Resealed nuclei restored transcription while inhibiting ESCRT-III activity accelerated cardiomyopathy. However, resealed nuclei were short-lived: they re-ruptured at twice the rate of resealing. A kinetic model predicted progressive accumulation of ruptured nuclei despite ongoing resealing. Consistently, a human LMNA-DCM heart contained numerous ruptured nuclei at disease presentation. These findings linked nuclear rupture to organ deterioration through global transcriptional deficiency and suggested rupture resealing as a critical modifier of nuclear rupture-associated conditions.

## MAIN

*LMNA*-related dilated cardiomyopathy (LMNA-DCM) is a predominantly adult-onset disease characterized by cardiac conduction defects and ventricular dilation that lead to life-threatening cardiac arrest or heart failure ^8^. LMNA-DCM is caused by certain heterozygous mutations in *LMNA*, which encodes for Lamin A/C, a critical structural component of the nuclear lamina ^9^. One prevailing model for the LMNA-DCM pathogenesis is that Lamin A/C deficiency or mutant expression weakens the nuclear envelope, ultimately causing disease. Supporting this, there are reports that LMNA-DCM patient hearts contain nuclei that appear to be ruptured ^10–12^. However, the magnitude and significance of nuclear rupture remain unknown. We previously reported that frequent nuclear rupture precedes cardiac structural and functional deterioration in a mouse model of LMNA-DCM ^7^, suggesting its potential pathogenic role. Nuclear rupture has also been recently reported in other pathological conditions, including cancer ^1,2,13^, neurodegenerative disease ^3–5^, myocardial infarction ^6^, other degenerative diseases associated with *LMNA* mutations ^14–18^, as well as certain physiological processes ^19,20^. Currently, whether nuclear rupture plays a pathogenic role in LMNA-DCM or any other rupture-associated diseases remains unclear.

One reason why nuclear rupture’s pathogenic role is unclear is that nuclear rupture is often transient due to endogenous nuclear envelope resealing activity ^1,21^. Consistently, nuclear rupture does not cause immediate cell death or impede cell cycle progression ^7,22^. A proposed mechanism for rupture sealing is the BANF1–ESCRT-III pathway, which normally orchestrates nuclear membrane reformation during mitosis ^1,21,23,24^. In addition, nuclear rupture can be resolved by the normal process of nuclear envelope reassembly during mitosis ^25–27^. While nuclear rupture sealing and resolution have been studied in cell culture, the fate of ruptured nuclei in disease contexts and in post-mitotic tissues remains unknown.

Another reason behind the unclear pathogenic role of nuclear rupture is the controversial molecular consequences of nuclear rupture. A prevailing hypothesis is that nuclear rupture causes DNA damage and a cytosolic DNA-driven immune reaction. While this hypothesis is supported in certain types of nuclear rupture, such as rupture of micronuclei ^28–30^, there are observations that do not fit in this model ^31,32^. For example, nuclear rupture in a mouse model of LMNA-DCM does not accompany immediate DNA damage or cGAS-STING cytosolic DNA-sensing activation ^7^. An impact on other nuclear processes has been overlooked despite substantial leakage of the nucleoplasm by nuclear rupture ^33^.

Here, we develop a novel reporter system that detects nuclear rupture and its resealing *in vivo*. Applying this to a mouse model of LMNA-DCM reveals ineffective rupture sealing and frequent re-rupturing, leading to accumulation of ruptured nuclei in the heart. We identify RNA polymerase II loss as a direct and pathogenic consequence of nuclear rupture in LMNA-DCM hearts.

## RESULTS

### Ruptured and resealed nuclei are both abundant in *Lmna^CKO^* hearts

To investigate the consequence of nuclear rupture and rupture sealing *in vivo*, we developed a dual reporter system that distinguishes nuclei presently ruptured from those resealed after initial rupture (**Fig. 1A**). This system employs co-expression of two different types of nuclear rupture reporters: a fluorescent protein fused with catalytically inactive cGAS (icGAS), a cytosolic DNA binding protein, and another fused with the nuclear localization signal (NLS). icGAS-based reporters are known to bind DNA at rupture sites and remain bound after rupture sealing ^1,21^, whereas NLS-based reporters are known to diffuse out of ruptured nuclei and reaccumulate back upon rupture sealing ^23,24^. Thus, combining the two markers would identify intact nuclei as icGAS^−^NLS^High^, ongoing ruptured nuclei as icGAS^+^NLS^Low^, and resealed nuclei as icGAS^+^NLS^High^ (**Fig. 1A**). We delivered GFP-icGAS and NLS-tdTomato to cardiomyocytes in mice via muscle-tropic adeno-associated virus 9 (MyoAAV) ^34^ or a Cre-inducible transgene, respectively.

**Figure 1.**
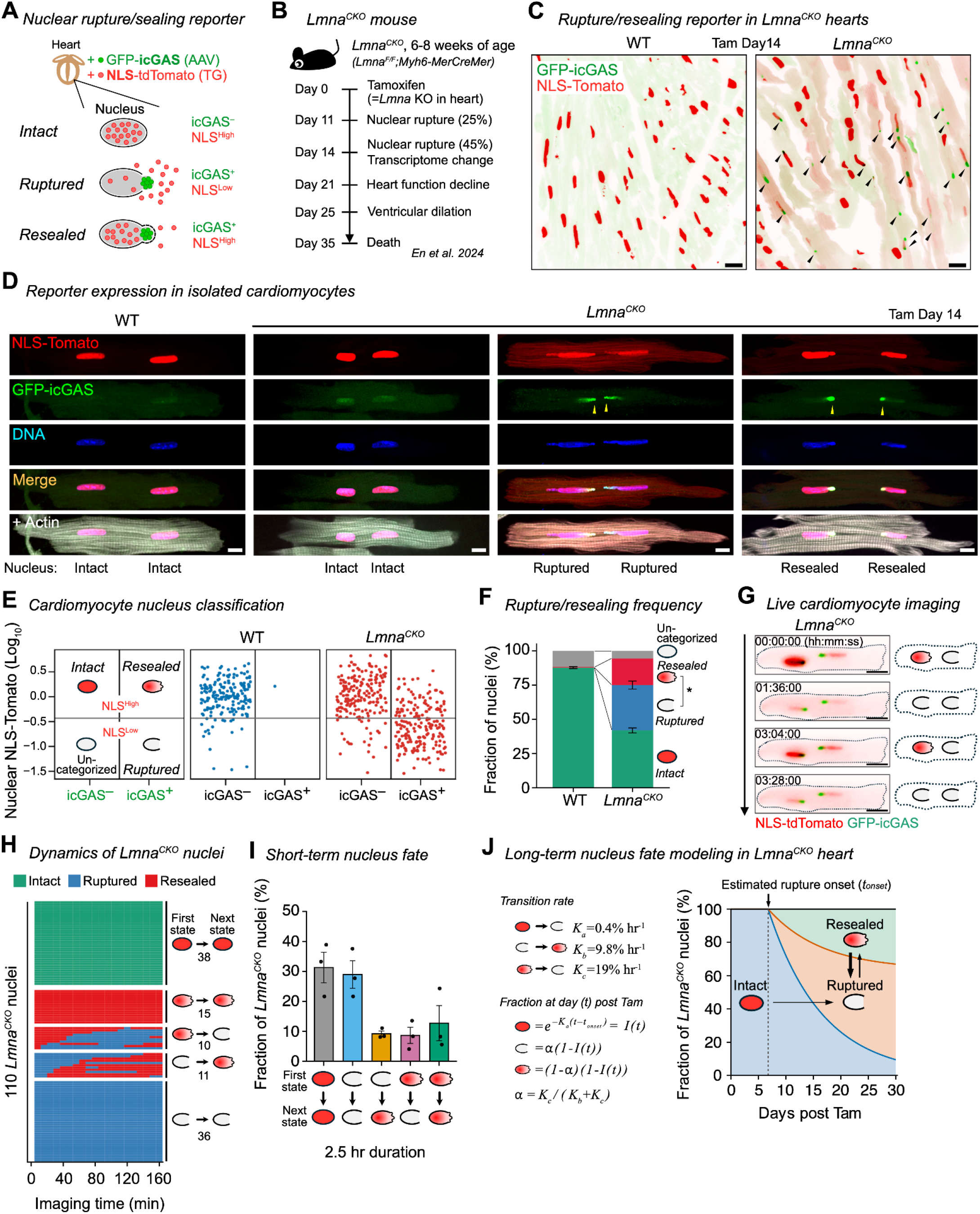
Dynamics of nuclear rupture and its resealing in *Lmna*^*CKO*^ hearts. **A)** GFP-icGAS and NLS-tdTomato reporters distinguish intact (icGAS^−^NLS^High^), ruptured (icGAS^+^NLS^Low^), and resealed (icGAS^+^NLS^High^) nuclei. GFP-icGAS and NLS-TdTomato are delivered to cardiomyocytes *in vivo* via AAV and a transgene (TG), respectively. **B)** Timeline of molecular, cellular, and pathological changes in *Lmna*^*CKO*^ mice after *Lmna* deletion induction in hearts. **C)** Heart sections expressing NLS-tdTomato and GFP-icGAS in wild-type (WT) and *Lmna*^*CKO*^ mice. Arrowheads: icGAS puncta at rupture sites. Scale bar: 20 μm. All data in Fig. 1 except 1J are derived from mice at day 14 post tamoxifen. **D)** Cardiomyocytes isolated from mice expressing NLS-tdTomato and GFP-icGAS. Each image shows one representative binucleated cardiomyocyte. Texts below images indicate nuclear states. Arrowheads: GFP-icGAS puncta at rupture sites. Scale bar: 5 μm. **E)** Classification of nuclei within cardiomyocytes by GFP-icGAS punctum (x-axis) and nuclear NLS-tdTomato intensity (y-axis). Horizontal line: mean minus one standard deviation of tdTomato intensity in icGAS^−^ nuclei. Data point: 188 WT nuclei and 390 *Lmna*^*CKO*^ nuclei (icGAS^+^ 204, icGAS^−^ 186) from 3 mice per genotype. See **Fig. S2** for individual replicates. **F)** Proportion of intact, ruptured, and resealed nuclei in cardiomyocytes. Asterisk: p-value <0.05 from parametric bootstrap from a binomial generalized linear mixed model (GLMM). N=3 mice per genotype. **G)** Representative time-lapse image of a live *Lmna*^*CKO*^ cardiomyocyte undergoing repeated nuclear rupture and resealing. Cells are isolated at day 14 post tamoxifen and used immediately for live imaging. Scale bar: 20 μm. **H)** Nuclear state dynamics of 110 nuclei in *Lmna*^*CKO*^ cardiomyocytes during 2.5-hr time-lapse imaging. Row: individual nuclei. Column: imaging time. Nuclei are grouped by state dynamics (right schematic). Number below schematic: nucleus count. Nuclei from three mice are shown. See **Fig. S3** for individual replicates. **I)** Fraction of nuclei undergoing different types of state transitions during 2.5-hr time-lapse imaging of *Lmna*^*CKO*^ cardiomyocytes. Mean fraction (bars) from three independent *Lmna*^*CKO*^ mice (point) and standard error range (error bar) are shown. **J)** Mathematical modeling of intact, ruptured, and resealed nuclear proportions over time post tamoxifen. Left: transition rates estimated from experimental data (top) and the kinetic models for proportions (bottom). Right, predicted proportions of intact, ruptured, and resealed nuclei based on the models and the observed proportions at day 14.

We used this dual reporter in mice with tamoxifen-inducible cardiomyocyte-specific *Lmna* deletion (*Lmna*^*F/F*^*;Myh6MerCreMer*; *Lmna*^*CKO*^), a mouse model of LMNA-DCM, and littermate control mice (*Lmna*^*+/+*^*;Myh6MerCreMer*) (**Fig. 1B**). In this model, *Lmna* is deleted in cardiomyocytes upon tamoxifen administration at 6–8 weeks of age, resulting in ~50% reduction in Lamin A/C protein abundance in cardiomyocytes due to Lamin A/C’s long half-life ^35^ (**Fig. S1A**). *Lmna*^*CKO*^ hearts develop nuclear rupture in 45% of nuclei at day 14 post tamoxifen ^7^, a time point of the main focus in the present study (**Fig. S1B**). This time point precedes functional and structural deterioration of the hearts, which begins day 21 ^7^.

The dual reporter system revealed abundant ruptured and resealed nuclei in *Lmna*^*CKO*^ hearts at day 14 post tamoxifen (**Fig. 1C; Fig. S1C**). In mutant hearts, we observed numerous nuclei marked with a GFP-icGAS punctum, some of which were associated with cytoplasmic NLS-tdTomato signals. GFP-icGAS puncta were essentially absent in wild-type control hearts. We next isolated cardiomyocytes from the hearts and fixed them immediately for high-resolution microscopy (**Fig. 1D; Fig. S1D**). GFP-icGAS puncta in *Lmna*^*CKO*^ cardiomyocytes co-localized with chromatin protruding from nuclear tips, consistent with previous reports ^7,36,37^. We observed two types of icGAS^+^ nuclei as predicted. One type of icGAS^+^ nuclei showed reduced NLS-tdTomato levels surrounded by cytoplasmic NLS-tdTomato signals, indicating ruptured nuclei (icGAS^+^NLS^Low^). Another type retained strong nuclear NLS-tdTomato, consistent with resealed nuclei (icGAS^+^NLS^High^). There were also intact nuclei with no GFP-icGAS puncta and strong NLS-tdTomato in *Lmna*^*CKO*^ cardiomyocytes (icGAS^−^NLS^High^). Individual *Lmna*^*CKO*^ cardiomyocytes contained intact, ruptured, or resealed nuclei, or any combinations of these states, including identical states, within a cell (of note, 90% of cardiomyocytes are binucleated in adult mice ^38^).

We identified the frequency of rupture and resealing events using classification based on dual reporter expression (**Fig. 1E; Fig. S2A**). In *Lmna*^*CKO*^ cardiomyocytes, 33% of nuclei were ruptured (icGAS^+^NLS^Low^), 19% resealed (icGAS^+^NLS^High^), 42% intact (icGAS^−^NLS^High^), and 6% uncategorized (icGAS^−^NLS^Low^), on average (**Fig. 1F**). Ruptured nuclei consistently outnumbered resealed nuclei across three independent mice (**Fig. S2B**). In wild-type cardiomyocytes, 88% of nuclei were intact, with virtually no ruptured or resealed nuclei, and 12% were uncategorized. In summary, ruptured and resealed nuclei together accounted for approximately half of all cardiomyocyte nuclei in *Lmna*^*CKO*^ hearts, with ruptured nuclei predominating.

### Frequent re-rupturing of nuclei in *Lmna*^*CKO*^ hearts

To understand how ruptured nuclei became abundant despite rupture sealing, we conducted a time-lapse imaging of cardiomyocytes expressing the dual reporter, freshly isolated from *Lmna*^*CKO*^ mice at day 14 post tamoxifen (**Fig. 1G; Fig. S3A; Movie 1**). During this experiment, cardiomyocyte contraction, which has little contribution to nuclear rupture ^37^, was inhibited by cytochalasin D to enable imaging and prevent hypercontraction-associated cell death. We successfully imaged 110 live cardiomyocytes from *Lmna*^*CKO*^ mice (n=3) (**Fig. S3B**).

The imaging revealed that ruptured nuclei were more likely to remain ruptured than resealed. Of 47 *Lmna*^*CKO*^ nuclei that had been ruptured at the beginning of imaging (icGAS^+^NLS^Low^), 77% (36 nuclei) remained ruptured, while 23% (11 nuclei) became resealed (**Fig. 1H, I; Fig. S3C**). We further found that resealed nuclei frequently re-ruptured. Of 25 *Lmna*^*CKO*^ nuclei that started with the resealed state (icGAS^+^NLS^High^), 60% (15 nuclei) remained resealed, while 40% (10 nuclei) re-ruptured (**Fig. 1H, I**). Several *Lmna*^*CKO*^ cardiomyocytes even underwent multiple rounds of resealing and re-rupturing (**Fig. S3D**). In contrast, *de novo* rupture was far less frequent than re-rupturing, as none of the intact *Lmna*^*CKO*^ nuclei (icGAS^−^NLS^High^) gained GFP-icGAS during the 2.5-hr imaging (**Fig. 1H**). This low frequency was consistent with the predicted rate of *de novo* rupture (0.42% per hour; **Methods**) based on our previous time course investigation, which showed that 25% of nuclei had ruptured by day 11 post tamoxifen and 45% by day 14, as indicated by chromatin protrusion from nuclear tip holes (**Fig. S1B**) ^7^.

We modeled long-term nuclear state dynamics using a first-order kinetic state-transition model using the observed rates of resealing (9.8% per hour), re-rupturing (19.2% per hour), and *de novo* rupture (0.42% per hour) (**Fig. 1J; Methods**). This model predicted that nuclear rupture began at day 7 post tamoxifen. Between day 14 and day 25, when cardiac structure and function severely deteriorate ^7^, ruptured nuclei are predicted to increase from 33% to 54%, resealed nuclei from 19% to 31%, and intact nuclei to decrease from 42% to 16%. These analyses suggested that more frequent re-rupturing than resealing drives progressive accumulation of ruptured nuclei in *Lmna*^*CKO*^ hearts.

### Global transcriptional deficiency in ruptured nuclei

Identification of intact, ruptured, and resealed nuclei in *Lmna*^*CKO*^ cardiomyocytes offered a unique opportunity to investigate molecular consequences of nuclear rupture. We hypothesized that nuclear rupture might cause reduction of transcription due to a loss of the nucleoplasm. To test this hypothesis, we isolated cardiomyocytes from *Lmna*^*CKO*^ and wild-type mice expressing the dual reporter at day 14 post tamoxifen and cultured them with bromouridine (BrU) for two hours to label nascent transcripts produced during the incubation (**Fig. 2A**). We observed that intact nuclei of wild-type cardiomyocytes exhibited robust BrU signals specifically in the nucleus, indicating strong transcriptional activity. Intact nuclei of *Lmna*^*CKO*^ cardiomyocytes similarly exhibited strong transcriptional activity. However, ruptured nuclei showed reduction of nascent transcript signals compared to intact nuclei. In contrast, resealed nuclei restored nascent transcript production. We next shortened the BrU labeling time to one hour to minimize labeling of transcripts from multiple nuclear states. This confirmed that nascent transcription was significantly reduced in ruptured nuclei and restored to the level of intact nuclei in resealed nuclei (**Fig. 2B**). Furthermore, BrU signals were positively correlated with nuclear NLS-TdTomato level in *Lmna*^*CKO*^ nuclei (R = 0.31) (**Fig. 2C; Fig. S4A**), suggesting a mechanistic link between transcriptional loss and nucleoplasmic loss in ruptured nuclei.

**Figure 2.**
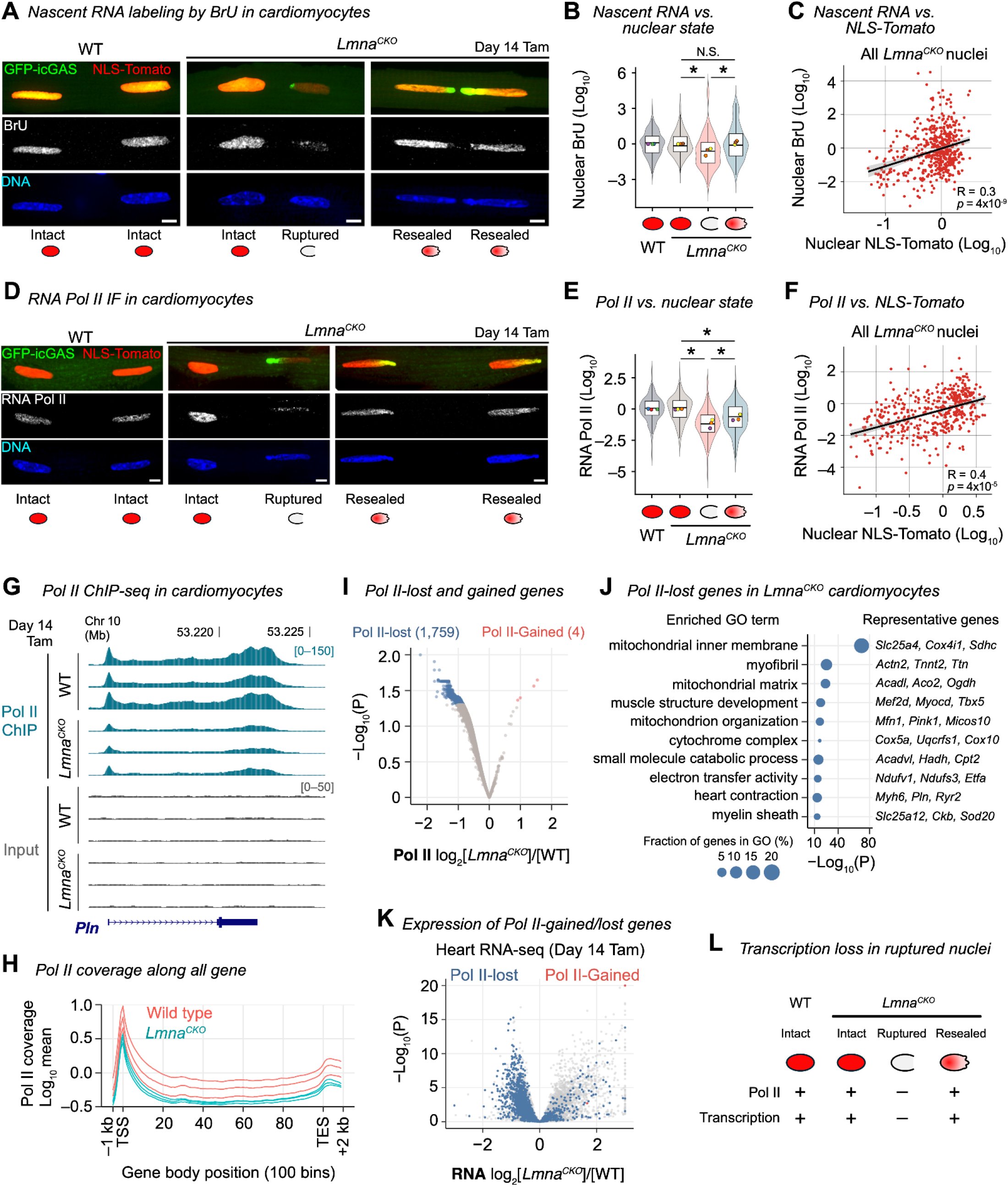
Loss of RNA polymerase II and transcriptional deficiency in ruptured nuclei. **A)** Immunofluorescence for bromouridine (BrU) after cardiomyocytes were cultured with BrU for 2 hr to label nascent RNAs. One binucleated cardiomyocyte per image, with nuclei zoomed in. Texts and icons below images indicate nuclear states. Scale bar: 5 μm. All data in Figure 2, except 2K, are from cardiomyocytes freshly isolated from mice at day 14 post tamoxifen. **B)** Nuclear BrU intensity by nuclear states after 1-hr BrU incubation. Density and box plots: BrU intensity distribution and interquartile range. Circles: mean intensity within biological replicates (color coded). Asterisks: P < 0.05 from *t*-tests on linear regression-estimated means with mouse-clustered standard errors. Underlying data: 244 intact nuclei from 4 WT mice, 275 intact, 111 ruptured, 104 resealed nuclei from 3 *Lmna*^*CKO*^ mice. **C)** Relationship between nuclear BrU intensity and nuclear NLS–tdTomato intensity in all types of nuclei in *Lmna*^*CKO*^ cardiomyocytes (519 nuclei from 3 mice). Line: simple linear regression fit with 95% confidence interval. R: Pearson correlation coefficient. P: *t*-test *p*-value on linear regression-estimated means with mouse-clustered standard errors. See **Fig. S4** for individual replicates. **D)** Immunofluorescence for RNA polymerase II (Pol II) in cardiomyocytes. Scale bar: 5 μm. **E)** Nuclear Pol II intensity by nuclear states. Underlying data: 285 intact nuclei from 3 WT mice, 180 intact, 133 ruptured, 136 resealed nuclei from 3 *Lmna*^*CKO*^ mice. Graph annotations and statistics as in **(B)**. **F)** Relationship between Pol II intensity and NLS–tdTomato intensity in all types of nuclei in *Lmna*^*CKO*^ cardiomyocytes (495 nuclei from 3 mice). Graph annotations and statistics as in **(C)**. **G)** Pol II ChIP-seq and input read coverage in WT and *Lmna*^*CKO*^ cardiomyocytes (3 mice per genotype). ChIP-seq signals are normalized to internal spike-in controls. **H)** Average Pol II ChIP-seq signals across 21,177 protein-coding genes. X-axis: 100 equally-spaced bins in gene bodies, 5 bins for 1 kb-upstream regions, and 10 bins for 2 kb-downstream regions. **I)** Statistical comparison of gene-body Pol II signals in *Lmna*^*CKO*^ versus WT cardiomyocytes for 11,942 Pol II-bound genes. Pol II-lost or gained genes are defined at *limma p*-value < 0.05. **J)** Ten most enriched Gene Ontology terms among the 1,759 Pol II-lost genes in *Lmna*^*CKO*^ cardiomyocytes, with three representative genes for each term. P: Metascape *p*-value. **K)** Gene expression state of Pol II-lost, Pol II-gained, and all other genes in *Lmna*^*CKO*^ (n=5) versus WT (n=7) hearts. P, DESeq2 *p*-value. RNA-seq data from En et al. 2024. **L)** Summary of Figure 2. Nuclear rupture causes transcriptional deficiency due to RNA Pol II loss.

### Loss of RNA Polymerase II in ruptured nuclei and its recovery by rupture sealing

We hypothesized that the observed transcriptional deficiency in ruptured nuclei was due to loss of RNA polymerases in the nucleoplasm. To test this hypothesis, we investigated the abundance of RNA Polymerase II (Pol II) using immunofluorescence in cardiomyocytes isolated from *Lmna*^*CKO*^ and wild-type mice at day 14 post tamoxifen. Pol II signals were abundant in intact nuclei in both wild-type and *Lmna*^*CKO*^ cardiomyocytes (**Fig. 2D**). In contrast, Pol II abundance was significantly reduced in ruptured nuclei, and then increased again in resealed nuclei, consistent with the changes in nascent transcription (**Fig. 2E**). However, Pol II abundance in resealed nuclei did not fully recover to the level of intact nuclei, mirroring the lower NLS-tdTomato signal in resealed nuclei compared with intact nuclei (**Fig. 1E**). Indeed, Pol II levels and nuclear NLS-tdTomato levels were strongly positively correlated among *Lmna*^*CKO*^ cardiomyocyte nuclei (R=0.41) (**Fig. 2F; Fig. S4B**). These data suggested that nuclear rupture caused leakage of soluble Pol II in *Lmna*^*CKO*^ cardiomyocytes.

Because soluble nucleoplasmic Pol II is the reservoir for transcriptionally-engaged Pol II, we investigated Ser5 phosphorylation (Ser5p) of Pol II C-terminal domain (CTD), a modification of initiating and early elongating Pol II (**Fig. S4C–F**), and CTD Ser2 phosphorylation (Ser2p), a modification of late elongating and terminating Pol II ^39^ (**Fig. S4G–J**). We observed that both Ser5p Pol II (**Fig. S4C, D**) and Ser2p Pol II (**Fig. S4G, H**) were reduced in abundance in ruptured nuclei. In resealed nuclei, Ser5p Pol II was restored to the level of intact nuclei, whereas Ser2p Pol II remained reduced, suggesting a slow recovery of late-stage elongating Pol II. Consistently, Ser5p Pol II levels were strongly positively correlated with nuclear NLS-tdTomato (R=0.43) (**Fig. S4E, F**), whereas Ser2p Pol II showed little correlation (R=0.05) (**Fig. S4I, J**). These data suggested that transcriptionally-engaged Pol II was reduced in ruptured nuclei in *Lmna*^*CKO*^ hearts.

### Genome-wide reduction of Pol II occupancy in *Lmna*^*CKO*^ cardiomyocytes

We next sought to identify genes that lost Pol II in ruptured nuclei. To this end, we performed ChIP-seq for Pol II in the whole *Lmna*^*CKO*^ cardiomyocyte population at day 14 post tamoxifen (**Fig. 2G**). We added a defined amount of human fibroblast chromatin to each ChIP reaction as an internal spike-in control to quantify Pol II occupancy change (**Fig. S5A**). We first confirmed the expected enrichment of Pol II signals at the 5’ end, gene body, and 3’ end in both *Lmna*^*CKO*^ and wild-type cardiomyocytes and in the spike-in human genome (**Fig. S5B, C**). We confirmed strong consistency of gene-body Pol II signals among biological replicates (**Fig. S5D**), but a clear separation by genotypes in principal component analysis (**Fig. S5E**). ChIP-seq profiles revealed that Pol II signals were reduced in gene bodies in *Lmna*^*CKO*^ cardiomyocytes, compared with the wild-type control (**Fig. 2G, H**). Of 11,942 protein-coding genes with measurable Pol II coverage, almost all (11,802, 99%) exhibited decreased gene-body Pol II coverage (log_2_ fold change < 0) (**Fig. 2I**). Among those, 1,759 genes showed the most consistent and strongest reduction (P < 0.05; **Table S1**). In contrast, only 4 genes showed increased Pol II at this threshold. These 1,759 Pol II-lost genes were dominated by Gene Ontology terms associated with cardiomyocyte homeostasis, function, and structure, and included such critical cardiac genes as *Cox4i1, Tnnt2, Myh6, Tbx5, Pln*, and *Ryr2* (**Fig. 2J**).

To test whether reduced Pol II occupancy was associated with decreased transcription, we assessed the expression of 1,759 Pol II-lost genes using whole-heart RNA-seq and single-nucleus (sn) RNA-seq datasets that we previously generated for wild-type and *Lmna*^*CKO*^ mice at day 14 post tamoxifen ^7^. A vast majority of the Pol II-reduced genes showed reduced expression (log_2_ fold change < 0) in *Lmna*^*CKO*^ hearts (75%; **Fig. 2K; Fig. S5F**) in *Lmna*^*CKO*^ pseudo-bulk cardiomyocyte population (65%; **Fig. S5G, H**). Together with nascent transcription analysis of individual nuclei, our data demonstrated that global transcriptional deficiency due to Pol II loss was the direct consequence of nuclear rupture in *Lmna*^*CKO*^ hearts (**Fig. 2L**).

### Lack of DNA damage response in ruptured and resealed nuclei

We next revisited the prevailing hypothesis that nuclear rupture causes DNA damage ^30^. In our previous work, we observed DNA damage accumulation in *Lmna*^*CKO*^ hearts at day 25 post tamoxifen, a terminal stage of the disease, but not at day 14 ^7^. We performed immunofluorescence for gamma-H2AX (phospho-S139 H2AX), a histone modification that marks DNA regions with double-strand breaks, in cardiomyocytes isolated at day 14. Low-level gamma-H2AX signals were present in intact nuclei of both wild-type and *Lmna*^*CKO*^ cardiomyocytes (**Fig. S6A**). Gamma-H2AX signals were slightly decreased, not increased, in ruptured nuclei compared to intact nuclei (P < 0.05; **Fig. S6B**). No significant difference was observed between ruptured and resealed nuclei or between intact and resealed nuclei. Accordingly, no significant relationship was observed between nuclear NLS-tdTomato levels and gamma-H2AX signal levels (**Fig. S6C**). These results further confirmed that nuclear rupture was not associated with increased DNA damage in *Lmna*^*CKO*^ cardiomyocytes.

### BANF1–ESCRT-III pathway proteins accumulate at nuclear rupture sites

We reasoned that identifying a rupture sealing mechanism would allow us to investigate the pathophysiological impact of nuclear rupture and its resealing in *Lmna*^*CKO*^ hearts. To date, nuclear membrane fusion driven by the BANF1–ESCRT-III pathway represents the prevailing model for rupture sealing ^1,21,23,24^. In this pathway, BANF1 binds cytoplasmic DNA and recruits LEMD2, which via CHMP7 recruits CHMP4B and other ESCRT-III machinery proteins. These proteins, together with VPS4 ATPase, catalyze membrane fusion without requiring nuclear lamins (**Fig. 3A**). To test whether this pathway operates in *Lmna*^*CKO*^ cardiomyocytes, we investigated the localization of BANF1, LEMD2, CHMP7, CHMP4B, and VPS4 in cardiomyocytes at day 14 post tamoxifen (**Fig. 3B**). Strikingly, all these five proteins were specifically localized to protruded chromatin in *Lmna*^*CKO*^ cardiomyocytes. In contrast, wild-type nuclei did not exhibit specific localization, except for LEMD2, which was weakly localized throughout the nuclear membrane. LEMD2 localization at protruded chromatin was noticeably distinct from that of other proteins (**Fig. S7A, B**): it appeared as sheets extending from intact regions of the nuclear membrane toward protruded chromatin tips, sometimes covering the entire chromatin protrusion. In contrast, BANF1, CHMP7, and VPS4 signals appeared as discrete foci.

**Figure 3.**
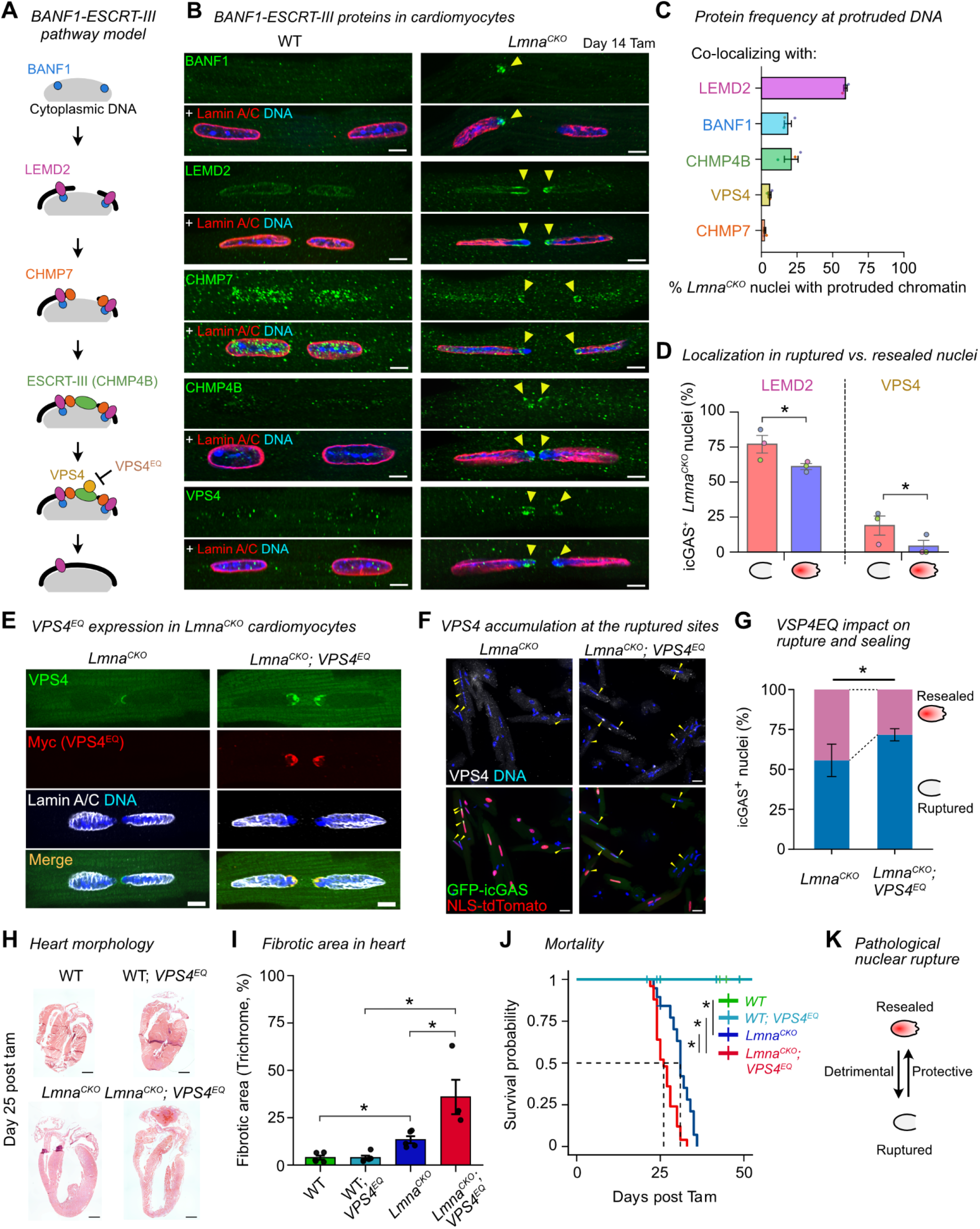
Rupture sealing by the BANF1–ESCRT-III pathway is cardioprotective. **A)** Model for the BANF1-ESCRT-III pathway driving nuclear membrane restoration on chromatin surface. **B)** Immunofluorescence for BANF1, LEMD2, CHMP7, CHMP4B, VPS4 and Lamin A/C in cardiomyocytes isolated at day 14 post tamoxifen. One binucleated cardiomyocyte per image, with nuclei zoomed in. Arrowheads: DNA protruded from rupture sites. Scale bar: 5 μm. **C)** Quantification of protruded chromatin colocalizing with BANF1-ESCRT-III pathway proteins. Graph shows mean percentage (bar) of biological replicates (points) with standard error ranges (error bars). Denominator: cardiomyocyte nuclei with protruded chromatin from 3 *Lmna*^*CKO*^ mice. Denominator nucleus count: LEMD2 n=156, BANF1 n=170, CHMP4B n=158, VPS4 n=156, and CHMP7 n=174. **D)** Quantification of ruptured or resealed nuclei with LEMD2 or VPS4 localization on protruded chromatin. Denominator: ruptured or resealed nuclei from 3 *Lmna*^*CKO*^ mice. Denominator nucleus count: LEMD2 n=171 nuclei (86 ruptured, 85 resealed), and VPS4 n=177 nuclei (119 ruptured, 58 resealed). Asterisk: *p*-value < 0.05 from Wald z-tests in binomial GLMMs estimating the probability of association. N=3 mice per genotype. **E)** Immunofluorescence for total VPS4, VPS4^EQ^ (Myc), and Lamin A/C in cardiomyocytes isolated at day 14 post tamoxifen. Scale bar: 5 μm. **F)** Immunofluorescence for total VPS4 in cardiomyocytes expressing GFP-icGAS and NLS-tdTomato at day 14 post tamoxifen. Arrowheads: icGAS puncta. Scale bar: 20 µm. **G)** Proportion of ruptured and resealed nuclei in *Lmna*^*CKO*^ (n=4) or *Lmna*^*CKO*^*;VPS4*^*EQ*^ (n=8) mice at day 14 post tamoxifen. Asterisk: *p*-value <0.05 from Wald z-tests in binomial GLMMs comparing the fraction of ruptured vs. resealed nuclei between genotypes. **H)** Gross morphology of hearts stained with Hematoxylin-Eosin at day 25 post tamoxifen. Scale bar: 1 mm. **I)** Fibrotic areas of myocardium positive for Masson’s Trichrome staining. Graph shows mean percentage (bar) of biological replicates (points) with standard error ranges (error bars). WT (n=5), WT;*VPS4*^*EQ*^ (n=5), *Lmna*^*CKO*^ (n=5), *Lmna*^*CKO*^*;VPS4*^*EQ*^ (n=4). Mean ± standard error. Asterisk: *p*-value <0.05 in one-way ANOVA with Tukey’s post-hoc test. **J)** Kaplan-Meier survival analysis. WT (n=16), WT;*VPS4*^*EQ*^ (n=23), *Lmna*^*CKO*^ (n=19), *Lmna*^*CKO*^*;VPS4*^*EQ*^ (n=25). Asterisk: *p*-value <0.05 in log-rank test. **K)** Summary of Figure 3. Nuclear rupture is detrimental to hearts while rupture sealing is cardioprotective.

The frequency at which BANF1–ESCRT-III proteins localized to protruded chromatin varied substantially. LEMD2 was most frequently observed at protruded chromatin (59%), followed by CHMP4B (21%), BANF1 (19%), VPS4 (6%), and CHMP7 (2%) (**Fig. 3C; Fig. S7C**). LEMD2 localization was frequent in both ruptured (77%) and resealed (61%) nuclei, with only a minor statistical difference (P=0.044; **Fig. 3D; Fig. S7D**). In contrast, VPS4 was much less frequent in resealed nuclei (4%) compared with ruptured nuclei (19%) (P < 0.05). These observations were consistent with the notion that LEMD2 remains in newly formed nuclear membranes ^40^, whereas the ESCRT-III-VPS4 complex dissociates from membranes after membrane remodeling ^41^. These observations supported the model that the BANF1–ESCRT-III pathway participated in rupture resealing in *Lmna*^*CKO*^ cardiomyocytes.

### No evidence that Lamin A/C or chromatin condensation promoted rupture resealing

We next investigated two other proposed mechanisms for rupture resealing. Nucleoplasmic Lamin A/C is known to relocalize to rupture sites and proposed to facilitate resealing ^42,43^. Although Lamin A/C remained in *Lmna*^*CKO*^ cardiomyocytes (**Fig. S1A**), it was largely absent from rupture sites (**Fig. 3B; Fig. S7B**), indicating that it did not participate in resealing. Other studies proposed that condensation of protruded chromatin acts as a diffusion barrier ^44^. However, there was no statistically significant difference in the proportion of condensed protruded chromatin between ruptured and resealed nuclei (37% vs. 33%; P>0.05) (**Fig. S8A, B**). Moreover, there was no relationship between condensation levels of protruded chromatin and the extent of NLS-tdTomato leakage (**Fig. S8C**). These observations suggested that neither nucleoplasmic Lamin A/C nor DNA condensation participated in rupture resealing in *Lmna*^*CKO*^ cardiomyocytes.

### Inhibiting ESCRT-III activity exacerbates nuclear rupture and accelerates DCM

To define the pathological contribution of nuclear rupture, we next inhibited the BANF1–ESCRT-III activity by expressing ATPase-defective dominant-negative VPS4-E228Q mutant (VPS4-EQ) ^45^. VPS4-EQ expression was driven by a Cre-inducible transgene, and thus was restricted to cardiomyocytes and initiated at the onset of *Lmna* deletion. At day 14 post tamoxifen, VPS4-EQ protein was specifically localized to protruded chromatin in *Lmna*^*CKO*^ cardiomyocytes, similar to endogenous VPS4B (**Fig. 3E**), but at much higher frequency (**Fig. 3F**). VPS4-EQ expression increased the fraction of ruptured nuclei by 1.3-fold, at the expense of resealed nuclei, in *Lmna*^*CKO*^*;VPS4*^*EQ*^ mice (**Fig. 3G**). The fraction of total icGAS^+^ nuclei (i.e. ruptured plus resealed) remained unchanged, confirming unaffected *de novo* rupture (**Fig. S9A**). Importantly, VPS4-EQ did not accumulate at other cellular membranes in which ESCRT-III might function (**Fig. 3F**), suggesting that VPS4-EQ specifically inhibited nuclear rupture sealing.

*Lmna*^*CKO*^*;VPS4*^*EQ*^ mice exhibited an accelerated development of dilated cardiomyopathy (DCM) compared with *Lmna*^*CKO*^ mice. *Lmna*^*CKO*^*;VPS4*^*EQ*^ hearts developed similar ventricular chamber dilation as in *Lmna*^*CKO*^ hearts (**Fig. 3H**) but exhibited more advanced fibrosis at day 25 post tamoxifen (**Fig. 3I; Fig. S9B**). *Lmna*^*CKO*^*;VPS4*^*EQ*^ mice died significantly earlier than *Lmna*^*CKO*^ mice (median survival 27 days in *Lmna*^*CKO*^*;VPS4*^*EQ*^ mice versus 31 days in *Lmna*^*CKO*^ mice, P=3×10^−4^; **Fig. 3J**). In contrast, *VPS4*^*EQ*^ single mutant mice did not show any chamber dilation or cardiac fibrosis or accelerated death, suggesting that the accelerated pathology in double mutants was due to specific perturbation of rupture sealing. These observations supported the hypothesis that nuclear rupture was pathological, whereas BANF1–ESCRT III-mediated rupture sealing was an endogenous cardioprotective mechanism in *Lmna*^*CKO*^ mice (**Fig. 3K**).

### Frequent nuclear rupture in a human *LMNA*-cardiomyopathy heart

Our data so far suggested that nuclear rupture accumulates despite ongoing resealing in Lamin A/C-deficient hearts. To assess whether similarly frequent nuclear rupture occurs in humans with *LMNA* mutations, we obtained a right ventricular biopsy from an individual carrying a heterozygous *LMNA* p.R541C (c.1621C>T) mutation (**Fig. S10A**). This heterozygous mutation was previously linked to DCM ^46,47^. The biopsy was taken after the subject underwent cardiac arrest followed by resuscitation at age 15. Using immunofluorescence for BANF1 as a rupture marker, we found that 19% of nuclei in the biopsy had a BANF1 punctum (**Fig. 4A–C**). BANF1 puncta were specifically localized at nuclear tips in cardiomyocytes, consistent with rupture-associated BANF1 observed in *Lmna*^*CKO*^ mouse hearts (**Fig. 4B**). BANF1 puncta were absent in a normal healthy human heart (**Fig. 4A, C**). They were also absent in an explant cardiomyopathy heart with a familial *ACTC1* (cardiac alpha-actin) pathogenic variant (**Fig. S10A**), excluding that BANF1 puncta were a general feature of cardiomyopathy. These data supported the hypothesis that ruptured nuclei accumulate in human hearts with *LMNA* mutations.

**Figure 4.**
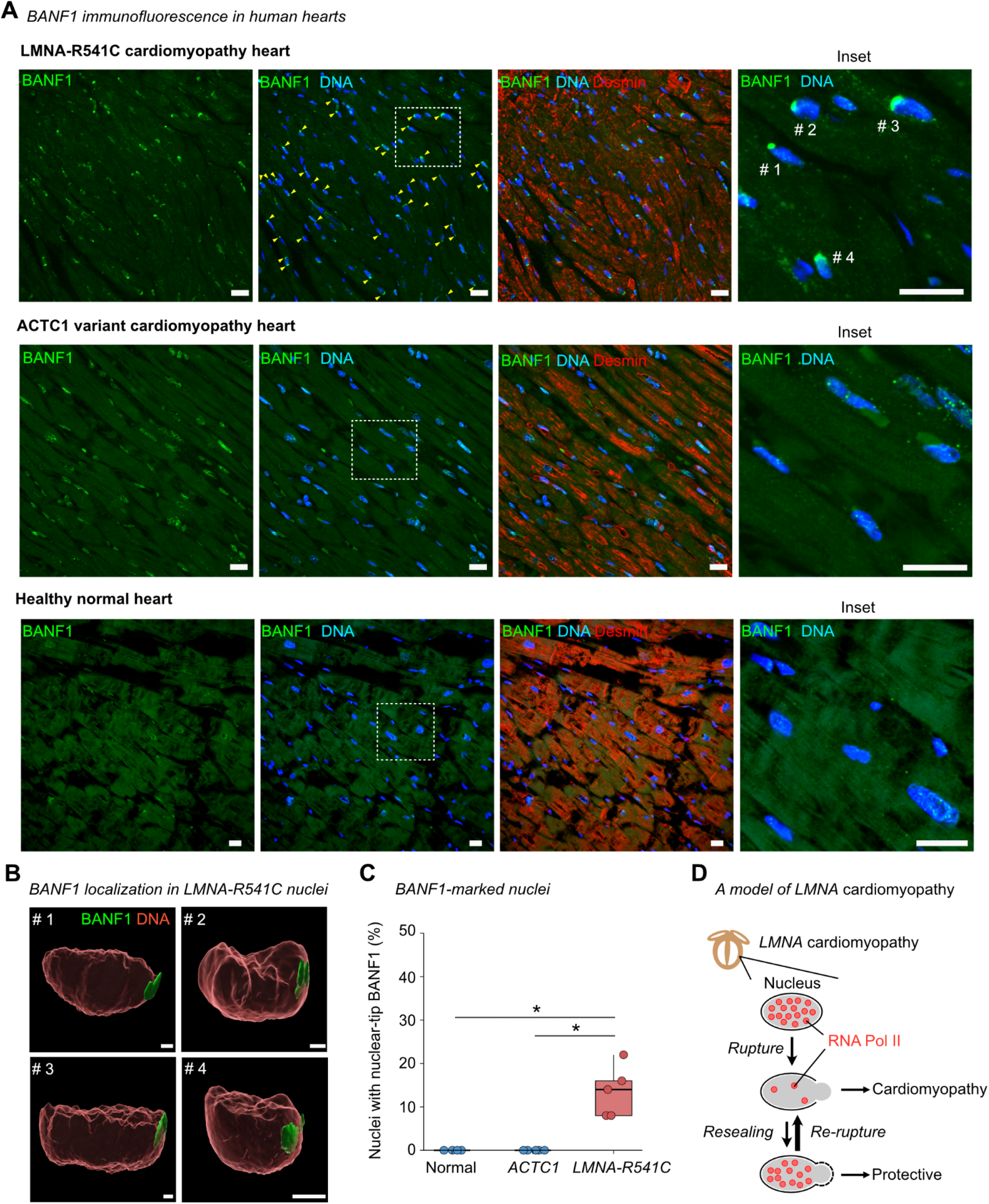
Frequent nuclear rupture in a human *LMNA*-cardiomyopathy heart. **A)** Immunofluorescence for BANF1 and Desmin in human hearts. Top: heart biopsy from an individual with a *LMNA* p.R541C heterozygous mutation. Middle: autopsy heart from an individual with a pathogenic *ACTC1* mutation. Bottom: apparently healthy autopsy heart. Inset: magnified areas enclosed by white squares. Scale bar: 20 μm. **B)** Three-dimensional reconstruction of BANF1 (green) and DNA (red) of nuclei (#1-4) in the *LMNA* p.R541C heart indicated in (**A**). Scale bar: 5 μm. **C)** Nuclei positive for nuclear-tip BANF1 per specimen field (normal heart, n=4 fields; *ACTC1* heart, n=5 fields, *LMNA*-R541C heart, n=5 fields). Box: interquartile range. Asterisk: *p*-value <0.05 in Kruskal–Wallis test with Holm-corrected Wilcoxon tests. **D)** Model. Nuclear rupture drives *LMNA*-cardiomyopathy due to Pol II loss. Ruptured nuclei accumulate in mutant hearts due to frequent re-rupture of resealed nuclei.

## DISCUSSION

This work identifies the consequence of nuclear envelope rupture and resealing *in vivo*. We show that RNA Pol II loss and the resulting global transcriptional deficiency are direct consequences of nuclear rupture in hearts (**Fig. 4D**). Our results suggest that inefficient rupture resealing and frequent re-rupturing drive the accumulation of ruptured nuclei in LMNA-DCM hearts. We predict that ruptured nuclei persist in post-mitotic adult hearts, because nuclei do not undergo mitotic nuclear envelope breakdown, a process that resolves nuclear rupture ^25–27^, and because nuclear rupture does not trigger immediate cell death ^7,22^. We propose that this persistence of transcriptionally defective nuclei drives cardiac tissue degeneration underlying cardiomyopathy. A similar mechanism may operate in other diseases involving nuclear rupture in post-mitotic cell types, such as in neurodegenerative diseases ^3–5^.

Our observation that re-rupturing was far more frequent than *de novo* rupture suggests that resealed nuclei are structurally weak. We propose that the lack of Lamin A/C re-recruitment to resealed sites, also observed in other cell types ^31^, contributes to this weakness. These findings suggest that stabilizing resealed nuclei is as important as preventing *de novo* rupture or increasing resealing frequencies when considering therapeutic strategies.

Our work provides several new insights into nuclear rupture resealing mechanisms. First, the BANF1–ESCRT-III pathway, an activity tied to mitosis-associated nuclear membrane reassembly, also operates in post-mitotic mammalian cells *in vivo*. Second, the sheet-like LEMD2 localization that we observed at rupture sites suggests that existing nuclear membranes extend outward *in cis* to seal ruptures, rather than that membranes are supplied *in trans* from other membrane systems for resealing. Third, our observation that VPS4 inhibition does not completely eliminate resealed nuclei suggests that mechanisms independent of the BANF1–ESCRT-III pathway also contribute to rupture sealing in cardiomyocytes.

While nuclear envelope rupture has been reported in various experimental conditions ^3,6,7,15^, its significance in human tissues has been obscure due to the lack of a liable marker. We show that BANF1 is a reliable marker of nuclear rupture and reveals frequent rupture in human LMNA-cardiomyopathy hearts. BANF1 has recently been used to detect nuclear rupture in various cancer tissues ^2^. We anticipate that our framework, combining BANF1-based rupture detection in human specimens with icGAS/NLS-based rupture and resealing detection in experimental systems, will advance understanding of the pathophysiological roles of nuclear rupture *in vivo*.

## ACKNOWLEDGEMENTS

We thank Dr. Harald Junge at the University of Minnesota and Dr. Katie Sue at Cincinnati Children’s Hospital Medical Center for VPS4-EQ transgenic mice, and Dr. Yixian Zheng at Carnegie Institution for *Lmna*-LoxP mice. We thank Ms. Jaime Reuss at Cincinnati Children’s for biopsy sample transfer. We thank Bio-Imaging and Analysis Facility, Genomics Sequencing Facility, Comparative Medicine Division for Animal Research, Integrated Pathology Research Facility, and Viral Vector Laboratory at Cincinnati Children’s. This work is supported by the National Institutes of Health (NIH) R01 HL176565 (K.I.) from the National Heart, Lung, And Blood Institute, NIH R21/R33 AG054770 (K.I.) from National Institute on Aging, Cincinnati Children’s Research Innovation and Pilot grant (K.I.), and Japan Society for the Promotion of Science Postdoctoral Fellowships for Research Abroad (W.K., A.E.). The content is solely the responsibility of the authors and does not necessarily represent the official views of the National Institutes of Health.

## AUTHOR CONTRIBUTION

Conceptualization, A.E. and K.I.;

Methodology, A.E., N.N., and K.I.;

Investigation, A.E., M.G., E.R., W.K., and K.I.;

Data Curation, A.E. and K.I;

Formal Analysis, A.E. and K.I.;

Writing – Original Draft, A.E. and K.I.;

Writing – Review & Editing, A.E. and K.I.;

Supervision, K.I.;

Project Administration, K.I.;

Resources, C.B., T.S., A.G., A.V.F., and K.N.W.;

Funding Acquisition, A.E. and K.I.

Validation, A.E. and K.I.;

Visualization, A.E. and K.I.;

## DATA AVAILABILITY

High-throughput sequencing data are available in Gene Expression Omnibus (GEO) website under GEO accession ID GSE325163.

## SUPPLEMENTAL TABLES AND MOVIES

**Table S1**

Gene body Pol II ChIP-seq coverage in wild-type and *Lmna*^*CKO*^ cardiomyocytes

**Table S2**

DNA oligonucleotide for genotype primer

**Movie 1**

Representative *Lmna*^*CKO*^ cardiomyocyte undergoing repeated re-rupture and resealing during 5-hr time-lapse imaging. Fluorescent signals are shown with (left) or without (right) bright-filed image.

## METHODS

### Mouse genetics

Experimental mice were housed in individually ventilated cages with up to 4 animals per cage. Animals were maintained under controlled environmental conditions: ambient temperature 22°C, relative humidity 30-80%, and a 14:12-h light–dark cycle. Animals had ad libitum access to irradiated standard chow and water.

Lmna-LoxP mice were provided by Dr. Yixian Zheng at Carnegie Institution and are available at the Jackson Laboratory (*Lmna*^*F/+*^; JAX stock no. 026284) ^48^. Myh6-MerCreMer mice (*Myh6MCM*^*Tg/+*^; JAX stock no. 005657) ^49^ and Rosa26-CAG-LSL-tdTomato mice (*R26*^*Tom/+*^; JAX stock no. 25106) ^50^ were obtained from the Jackson Laboratory. Rosa26-CAG-LSL-Vps4EQ mice (*R26*^*VPS4EQ/+*^) ^45^ were provided by Dr. Harald Junge at University of Minnesota Twin Cities and Dr. Katie Sue at Cincinnati Children’s Hospital Medical Center.

For cardiomyocyte-specific *Lmna* knockout experiments, we used *Lmna*^*F/F*^*;Myh6MCM*^*Tg/0*^ (*Lmna*^*CKO*^) and *Lmna*^*+/+*^*;Myh6MCM*^*Tg/0*^ (wild-type control) mice. For the analysis of nuclear tdTomato intensity, we used *Lmna*^*F/F*^*;Myh6MCM*^*Tg/0*^*;R26*^*Tom/+*^ and *Lmna*^*+/+*^*;Myh6MCM*^*Tg/0*^*;R26*^*Tom/+*^. For dominant negative VPS4EQ experiments, we used *Lmna*^*F/F*^*;R26*^*+/+*^*;Myh6MCM*^*Tg/0*^, *Lmna*^*+/+*^*;R26*^*+/+*^*;Myh6MCM*^*Tg/0*^, *Lmna*^*F/F*^*;R26*^*VPS4EQ/+*^*;Myh6MCM*^*Tg/0*^, and *Lmna*^*+/+*^*;Rosa26*^*VPS4EQ/+*^*;Myh6MCM*^*Tg/0*^ mice. The *Lmna*^*F/+*^ allele and *R26*^*VPS4EQ/+*^ allele were genotyped by PCR using the primers KI279 and KI280 or MEG036, MEG037, and MEG038 (**Table S2**). The *Myh6MCM*^*Tg/+*^ allele and *R26*^*Tom/+*^ allele were genotyped by Transnetyx, Inc using a probe pair of Esr-Cre TG and Chr19-2 WT and a probe pair of tdRFP and ROSA WT, respectively. All parental mice to experimental mice have been crossed with CD1 outbred mice (Charles River Laboratories) at least twice to ensure the robustness of the phenotypes independent of particular genetic backgrounds. Both sexes were used in experiments.

To induce Cre-mediated recombination, all experimental mice were administered with tamoxifen (Sigma, T5648) at 6–8 weeks of age via intraperitoneal injections (100 μL of 4 mg/mL solution per day for 4 consecutive days) dissolved in corn oil (Sigma, C8267).

All mouse experiments were approved by the Institutional Animal Care and Use Committee (IACUC) at Cincinnati Children’s Hospital under IACUC protocol 2024-0047. All procedures were performed in compliance with institutional and governmental regulations under PHS Animal Welfare Assurance number D16–00068 (Cincinnati Children’s).

### Human cardiac tissues

Human LMNA-R541C samples were provided as de-identified specimens under Cincinnati Children’s Hospital Institutional Review Board (IRB) protocol # 2020-0390. The use of human autopsy specimens provided by CVPath Institute was approved by the CVPath Institute Institutional Review Board.

### Cell culture

We used AAVpro 293T cells (TaKaRa, 632273) for MyoAAV production and BJ-5ta cells (human hTERT-immortalized foreskin fibroblast; ATCC, CRL-4001) for ChIP-seq spike-in controls. Both cell lines were cultured in high-glucose DMEM (Gibco, 11965-092) supplemented with 9% fetal bovine serum (FBS) and 1:100 penicillin–streptomycin (Invitrogen, 105727) at 37°C in 5% CO_2_.

### MyoAAV preparation and treatment

Preparation and administration of icGAS-GFP MyoAAV (pAAV:cTnT::GFP-icGAS) are detailed in our previous study ^7^. Briefly, the icGAS-GFP expression vector was generated by fusing icGAS cDNA (human cGAS with E225A/D227A substitutions) with EGFP cDNA and inserting this into the pAAV:cTNT vector (gift from William Pu; Addgene plasmid # 69915). MyoAAV was produced by co-transfecting the icGAS vector with the pHelper vector (GenBank: AF369965.1) and the pRep/Cap 1A-MYO capsid vector ^34^ into AAVpro 293T cells (TaKaRa, 632273). AAV was purified using a published protocol ^51^ which is also described in our previous paper ^7^. The viral genome copy numbers (vg) were estimated by qPCR ^7^. MyoAAV particles were administered to mice through retro-orbital venous sinus injection at a dose of 1–2×10^11^ vg per mouse at 4 weeks of age, which was 2 weeks prior to tamoxifen.

### Survival analysis

Kaplan-Meier survival analyses were performed using the *survival* package and visualized with the *survminer* package in R. Log rank tests were performed using the *survdiff* function in the *survival* package.

### Histological staining

Hearts were perfused with 100 mM KCl in PBS and fixed in 10% formalin (Fisherbrand, 245–685). Fixed hearts were embedded in paraffin and sectioned to a thickness of 5 μm. Heart sections were deparaffinized using xylene and ethanol and stained with hematoxylin and eosin (H&E) or a Masson’s Trichrome Kit (Thomas Scientific LLC, KTMTRPT) according to the manufacturer’s protocol. Stained heart images were obtained on Nikon NiE microscope for H&E stained sections or Olympus BX51 microscope for Trichrome stained sections. The proportion of Trichrome-positive areas was quantified from 3–5 randomly selected, non-overlapped images per mouse, and the mean proportion per mouse was used for plots and statistical analysis (see Statistical Analysis).

### Cardiomyocyte isolation

Mice were administered with heparin (100 Units, NDC 25021–400-10), anesthetized with isoflurane, and euthanized by cervical dislocation. Hearts were perfused with 100 mM KCl in PBS and moved to Potassium Buffer (4.65 mM 2,3-butanedione monoxime, 0.5 mM EGTA, 11.1 mM glucose, 40.2 mM KCl, 16.2 mM KH2PO4, 70 mM L-Asp potassium salt, 10 mM Na-pyruvate, 10 mM taurine, 10.1 mM HEPES, pH 7.4). Excised hearts were cannulated to the Langendorff retrograde perfusion system through the aorta and perfused with prewarmed Digestion Buffer (1 mg/ml Collagenase Type II, Worthington, LS004177, 10 mM glucose, 5 mM HEPES, 5.4 mM KCl, 1.2 mM MgCl2, 150 mM NaCl and 2 mM sodium pyruvate, 10 mM taurine and 12 mM 2,3-butanedione monoxime, pH 7.35) for 20 min at 37°C. For immunostaining and live imaging analysis, cardiomyocytes were dissociated from digested hearts by gently mincing in a Potassium Buffer supplemented with 5 mg/mL BSA. Cardiomyocyte suspension was filtered through a 240 μm mesh. Cardiomyocytes settled to the bottom of conical tubes by gravity were used in experiments.

For chromatin immunoprecipitation, digested hearts were perfused with 10 mL of 1% paraformaldehyde (PFA; Electron Microscopy Slides, #15710) in PBS prior to cell dissociation. Cardiomyocytes were then dissociated by gently triturating in 1% PFA in PBS. Cell suspension was filtrated through a 240 μm mesh and incubated with 1% PFA for a total 10 min (from perfusion to this step) and quenched by 125 mM glycine. Cardiomyocytes were sedimented at the bottom of conical tubes by gravity, and the resulting cell pellet was further purified in PBS using gravity. Fixed cardiomyocytes were snap-frozen by liquid nitrogen and stored at –80°C until use.

### Nascent transcript labeling using bromouridine

*Lmna*^*CKO*^ cardiomyocytes isolated at 2 weeks post tamoxifen were resuspended in Potassium Buffer supplemented with 25 µM Cytochalasin D to halt cardiomyocyte contraction. Calcium concentration was gradually increased from 0.5 mM to the final concentration of 2.0 mM by adding CaCl_2_ every 3 min. Cells were resuspended in Medium 199 (Gibco, 11150-059) supplemented with 1x Insulin-Transferrin-Selenium-X (Gibco, 51500-056), 1:100 penicillin–streptomycin (Invitrogen, 105727), 25 µM Cytochalasin D, and 20 mM HEPES (pH 7.4), and plated onto coverslips coated with laminin (Gibco, 23017015) on dishes. Cells were incubated with 2 mM bromouridine (Sigma Aldrich, 850187) in fresh supplemented Medium 199 for either 1 hr or 2 hr at 37°C in 5% CO_2_. We used 1-hr incubation for quantification of nascent RNA and 2-hr incubation for visualization. After labeling, cells were fixed with 2% paraformaldehyde in PHEM buffer (60 mM PIPES pH7.5, 25 mM HEPES pH7.5, 10 mM EGTA, 4 mM MgSO_4_) at 37°C for 10 min and processed for immunostaining.

### Immunofluorescence in isolated cardiomyocytes

Isolated cardiomyocytes were fixed in 2% PFA in PHEM buffer (60 mM PIPES pH7.5, 25 mM HEPES pH7.5, 10 mM EGTA, 4 mM MgSO4) for 10 min at 37°C, then washed with PBS and attached to coverslips with poly-L-lysine (Sigma Aldrich, P4832). Prior to blocking and permeabilization, following pretreatment steps were included in some staining experiments. For staining of DNA-binding proteins (BANF1 and gamma-H2AX), cells were subjected to heat-induced antigen retrieval in 10 mM citrate buffer (pH 6.0) at 60 °C for 15 min to improve antibody accessibility to condensed chromatin at nuclear rupture sites. For co-staining with mouse anti-VPS4 and mouse anti-c-Myc antibodies in cardiomyocytes expressing NLS-tdTomato and GFP-icGAS, endogenous fluorescent signals were bleached in alkaline solution (1% NaOH, 0.9% NaCl) for 1 hr at room temperature ^52^. In all experiments, cells were blocked and permeabilized in a buffer containing 5% normal donkey serum (Jackson ImmunoResearch, 017–000-121), 1% non-fat milk, and 0.1% Triton X-100 in PBS for 1 h at 37°C. Permeabilized cells were incubated overnight at 4°C with the following primary antibodies. Rabbit anti-gamma-H2AX antibody (Cell signaling, 9718); mouse anti-BrU/BrdU antibody (BD Biosciences, 555627); rabbit anti-RNA Pol II N-terminal domain (NTD) antibody (Cell signaling, 14958); rat anti-RNA Pol II CTD phospho-Ser5 antibody (Chromotek, AB_2631404); rat anti-RNA Pol II CTD phospho-Ser2 antibody (Millipore Sigma, 04-1571); mouse anti-Lamin A/C antibody (Santa Cruz, sc-376248); rabbit anti-Lamin A antibody (Abcam, ab26300); rabbit anti-PCM1 antibody (Sigma, HPA023370); mouse anti-BANF1 antibody (Abnova, H00008815-M01); rabbit anti-LEMD2 antibody (Millipore Sigma, HPA017340); rabbit anti-CHMP4B antibody (Proteintech, 13683-1-AP); rabbit anti-CHMP7 antibody (Proteintech, 16424-1-AP); mouse anti-VPS4 antibody (Santa Cruz, sc-133122); and mouse anti-c-Myc antibody (Invitrogen, 13-2500). Cells were washed and incubated with Alexa fluorophore-conjugated secondary antibodies and Alexa Fluor Plus 647-conjugated Phalloidin for 1 h at 37°C. For co-staining of mouse anti-VPS4 and mouse anti-c-Myc antibodies, subclass-specific nanobodies were used for detection (anti-mouse IgG2a nanobody, Proteintech, smsG2aCL488-1; anti-mouse IgG1 nanobody, Proteintech, smsG1CL647-1). Cells were counterstained with DAPI (4’,6-diamidino-2-phenylindole) and then mounted with ProLong Glass Antifade Mountant (Thermo Fisher P36980). For combinatorial detection with the native fluorescent reporters NLS-tdTomato and GFP-icGAS, immunostained signals (gamma-H2AX, BrU, RNA Pol II NTD, RNA Pol II CTD pSer5, RNA Pol II CTD pSer2, LEMD2, and VPS4) were detected using Alexa Fluor Plus 647–conjugated secondary antibodies.

Immunofluorescence signals and native fluorescent signals (tdTomato and icGAS) were acquired using a Yokogawa CSU-W1 Sora spinning-disk confocal microscope with a 40x or 60x objective using a 0.2-µm Z-step size. We imaged at least 3 areas in one sample (coverslip), each of which roughly contain ~50 cardiomyocytes. Because isolated cells were randomly distributed on the coverslip relative to their original tissue location, the images reflect average cell states in the heart.

### Nuclear state identification in fixed cardiomyocytes using icGAS/NLS reporters

All cardiomyocytes maintaining the rod shape and containing two or more nuclei in each image were included in analyses. To determine icGAS^+^ and icGAS^−^ nuclei, we determined whether a distinct GFP-icGAS punctum was present at the nuclear tip for each nucleus. We quantified nuclear NLS-tdTomato levels using either Fiji image analysis software ^53^ for quantifying nucleus states or using Imaris software (Oxford Instruments) for quantifying immunofluorescence signals within different nucleus states (see next section), using essentially the same analytical approach. In Fiji, we first generated maximum projection intensity projections of z-stack images, defined a two-dimensional region of interest (ROI) for each nucleus (~25–75% of the nuclear area), and calculated the log_10_–transformed mean NLS-tdTomato pixel intensity within each ROI (*NucTom*). In Imaris, we first generated a three-dimensional ROI for each nucleus using DAPI signal segmentation and calculated the log_10_–transformed mean NLS-tdTomato pixel intensity within each ROI (*NucTom*). The downstream steps were identical between the two approaches. We first normalized each nucleus’s tdTomato intensity to the mean tdTomato intensity of icGAS^−^ nuclei within the same mouse (i.e. same coverslip) (*NormNucTom*). We then defined the threshold to distinguish resealed and ruptured nuclei as the mean minus one standard deviation of the normalized tdTomato intensity of icGAS^−^ nuclei in each mouse (*Threshold*). For each mouse, icGAS^+^ nuclei with normalized tdTomato intensity less than or equal to this threshold were categorized as ruptured (icGAS^+^NLS^Low^), whereas those greater than this threshold were categorized as resealed (icGAS^+^NLS^High^). icGAS^−^ nuclei with normalized tdTomato intensity greater than this threshold were categorized as intact (icGAS^−^NLS^High^), whereas those less than or equal to this threshold were categorized as uncategorized (icGAS^−^NLS^Low^). Mathematically, for nucleus *i* in mouse *j*:

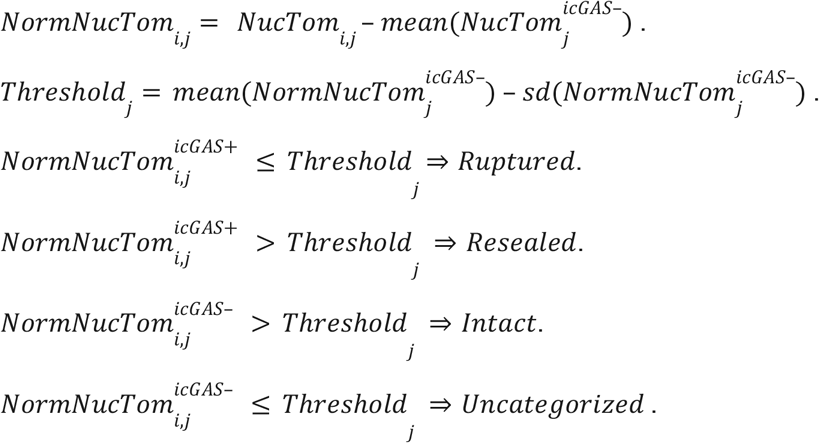

Importantly, we used nuclear NLS-tdTomato intensity, instead of a nuclear-to-cytoplasmic ratio, for nuclear state analysis because cytoplasmic signals are influenced by the number of ruptured or resealed nuclei per cell. We previously established that nuclear NLS-tdTomato intensity is highly consistent across intact nuclei in adult post-mitotic cardiomyocytes, due to the absence of mitotic nuclear envelope breakdown ^7^. See the *Statistical analysis* section for statistics.

### Quantification of ever-ruptured nuclei using PCM1 staining

We used DNA protrusion at nuclear tips accompanied by local PCM1 signal depletion to quantify ever-ruptured nuclei (i.e. ruptured and resealed nuclei) at day 11 or day 14 post tamoxifen. Of note, protruded DNA at PCM1-loss sites was almost always colocalized with GFP-icGAS in *Lmna*^*CKO*^ cardiomyocytes and thus serves as a reliable marker of ever-ruptured nuclei ^7^. We reanalyzed PCM1 immunofluorescence data from isolated cardiomyocytes at day 11 or day 14 post tamoxifen that we previously reported ^7^. This reanalysis was necessary because the prior study quantified cardiomyocytes containing ruptured nuclei, not the number of ruptured nuclei.

### Quantification of immunofluorescence signals in nuclei

Quantification of immunofluorescence signals was performed using Imaris image analysis software (Oxford Instruments). For all analyses, nuclei were segmented based on DAPI signals.

For quantification of BrU, Pol II, Pol II Ser2p, Pol II Ser5p, and gamma-H2AX, we first obtained the sum (BrU, Pol II, Pol II Ser2p, and Pol II Ser5p) or mean (gamma-H2AX) of signal intensities within each segmented nucleus and transformed it into the log_10_ scale (*NucFL*). The log_10_ signals were normalized by the mean and standard deviation of the signals within intact nuclei (icGAS^−^NLS^High^). Mathematically, normalized nuclear fluorescence intensity (*NormNucFL*) for nucleus *i* in mouse *j* was:

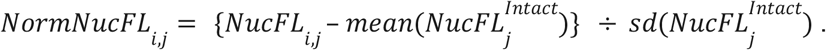

Normalized signals were used in data plots. See the *Statistical analysis* section for statistics.

### Immunofluorescence signal analysis at protruded chromatin

Immunofluorescence signals for BANF1, LEMD2, CHMP7, CHMP4B, and VPS4 at protruded chromatin were analyzed in three dimensions using Imaris. Nucleus regions including protruded chromatin were defined by DAPI signals. Separately, the nuclear envelope surface was defined by Lamin A/C for each nucleus, and the ruptured surface was identified as the local opening of the nuclear envelope at the nuclear tip. For each nucleus, the nucleus region was separated into the main nucleus compartment and the protruded chromatin at the ruptured surface using the *cut* function in Imaris. For each nucleus, we measured the mean immunofluorescence intensity in the main nucleus compartment (“a”) and the protruded chromatin part (“b”). The log_2_ ratio (b/a) was used to quantify relative protein abundance at protruded chromatin (*RelAbs*). For statistical analysis, we estimated a null distribution by computing the log_2_ ratio of the main nucleus signal relative to the mean main nucleus signal across all nuclei (*RelAbs*^*Null*^). Protruded chromatin regions with log_2_(b/a) greater than the mean plus two standard deviations of this null distribution (empirical *p* < 0.05) and greater than 0.585 (log_2_ 1.5) were considered positive for factor localization. Mathematically, for nucleus *i* in mouse *j*:

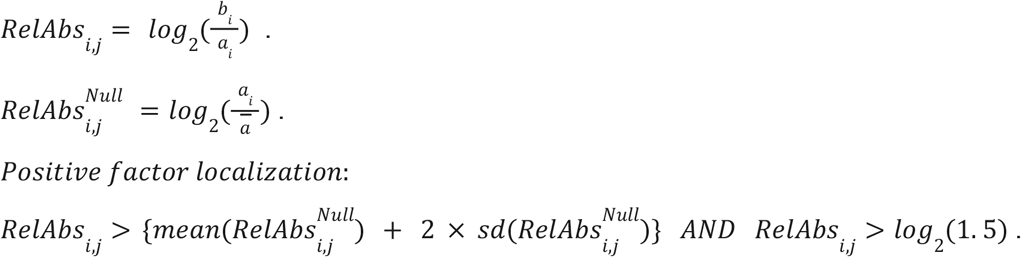

For signal comparisons between ruptured and resealed nuclei, including DAPI intensities, we used the same framework except that the protruded chromatin region was defined by the icGAS-positive area, and the main nucleus compartment was obtained by geometrically subtracting this region from the total nucleus volume. See the *Statistical analysis* section for statistics.

### Time-lapse live imaging of isolated mouse cardiomyocytes

Cardiomyocytes isolated from *Lmna*^*CKO*^ mice at 2 weeks post tamoxifen were resuspended in Potassium Buffer supplemented with 25 µM Cytochalasin D to maintain cell viability and minimize motion artifacts during imaging. Calcium concentration was gradually increased from 0.5 mM to the final concentration of 2.0 mM by adding CaCl_2_ every 3 min. Cells were resuspended in Medium 199 (Gibco, 11150-059) supplemented with 1x Insulin-Transferrin-Selenium-X (Gibco, 51500-056), 1:100 penicillin–streptomycin (Invitrogen, 105727), 25 µM Cytochalasin D, and 20 mM HEPES (pH 7.4), and plated onto glass-bottom dishes (MatTek D35C4-20-1.5-N) coated with laminin (Gibco, 23017015). Dishes were placed in a stage-top humidified incubator maintained at 37°C and 5% CO_2_ (Tokai Hit) in a Nikon TiE inverted microscope equipped with a SpectraX light source. Images were acquired at four positions in the dish every 8 minutes for a total of 5 hours using a 10x objective. GFP-icGAS and NLS-tdTomato were sequentially imaged using 470-nm and 555-nm excitation with optimized filter settings to minimize switching time during time-lapse acquisition. (**Movie 1**).

### Time-lapse image analysis

The analysis of icGAS-GFP and NLS-tdTomato signals was performed on cardiomyocytes that were morphologically normal, survived for a 2.5-hour duration, and contained two or more nuclei, using Fiji image analysis software. We chose this 2.5-hour analysis duration from the total 5-hour imaging time because this duration allowed analysis of abundant healthy viable cells. icGAS^+^ and icGAS^−^ nuclei were identified as described above in the analysis of fixed cells (*Nuclear state identification in fixed cardiomyocytes*). For NLS-tdTomato analysis, we first created an ROI within the nuclear area for each nucleus for each time point, obtained mean nuclear tdTomato intensity within the ROI, and log_10_-transformed it (*NucTom*). We normalized each nuclear tdTomato intensity to the mean nuclear tdTomato intensity across all icGAS^−^ nuclei within that time point (*NormNucTom*). This value was smoothed by taking a median across three consecutive frames (*t*−1, *t, t* + 1) (*SmoNormNucTom*). The resulting smoothed normalized tdTomato intensity was visualized in line plots for nuclear tdTomato signals. The threshold for distinguishing sealed and ruptured nuclei was defined as the mean minus one standard deviation of the smoothed normalized nuclear tdTomato intensity in icGAS^−^ nuclei for each time point (*Threshold*). For nuclear state classification, nuclei with normalized tdTomato signals greater than the threshold above were classified as intact if icGAS^−^ or resealed if icGAS^+^, whereas nuclei with values less than or equal to the threshold were classified as ruptured if icGAS^+^. This classification was visualized in the heatmap. Mathematically, for nucleus *i* in mouse *j* at time *t*:

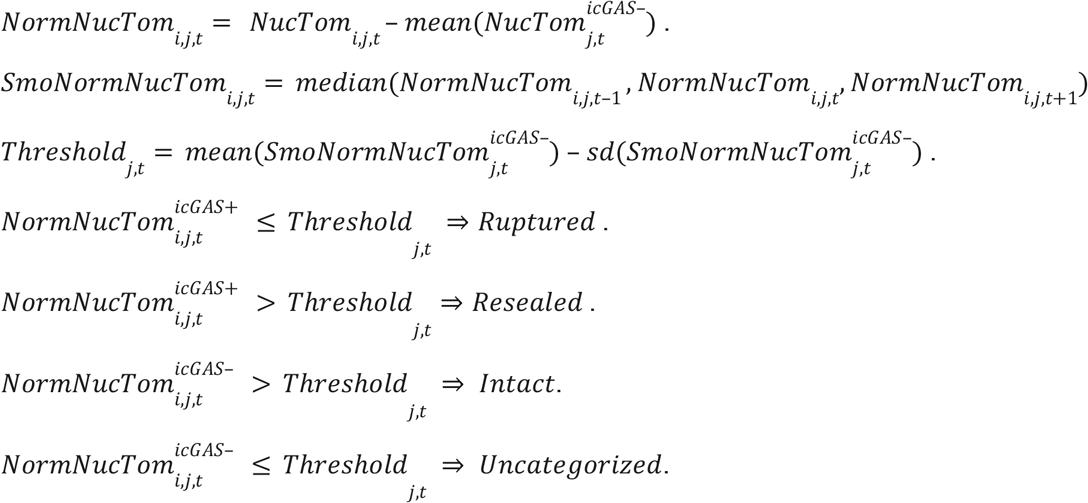

To visualize nucleoplasmic and cytoplasmic tdTomato intensity side-by-side in individual cardiomyocytes, a second ROI of identical size was placed in the cytoplasm adjacent to each nucleus to quantify cytoplasmic tdTomato intensity. Because the two nuclei reside within a shared cytoplasm in binucleated cardiomyocytes, cytoplasmic intensities measured from the two ROIs were averaged to generate a single cytoplasmic trace for each cell. Raw nuclear (*NucTom*) and cytoplasmic tdTomato intensity (*CytTom*) traces were smoothed using a three-frame running median filter (*t*−1, *t, t* + 1) (*SmoNucTom*; *SmoCytTom*), mean-centered (*CentSmoNucTom*; *CentSmoCytTom*;), and the nuclear trace was scaled by 0.1 for visualization. Mathematically, for nucleus *i* at time *t*, with *T* indicating the final time point:

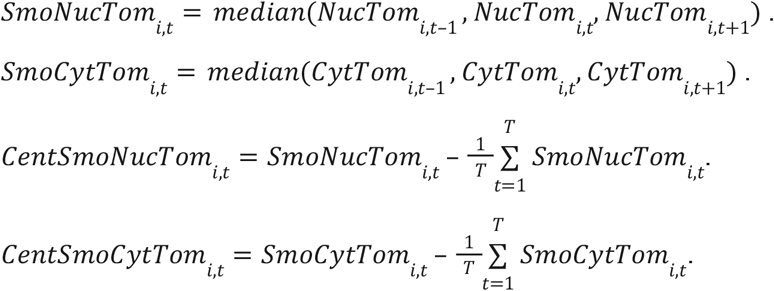

### Modeling of nuclear rupture and resealing dynamics

We constructed a kinetic model with intact, ruptured, and resealed nuclear states. Transitions were modeled using first-order kinetics, in which we assumed each nucleus had a constant probability per unit time of transitioning to another state. Consequently, the total transition rate is assumed proportional to the fraction of nuclei currently in that state. Therefore, the rate of change in intact (*I*), ruptured (*R*), and resealed (*S*) nucleus fractions could be estimated as:

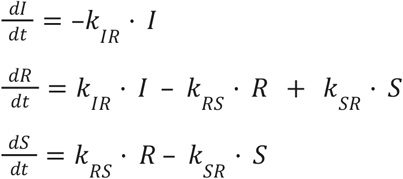

where *k*_*IR*_ is the *de novo* rupture rate, *k*_*RS*_ the resealing rate, and *k*_*SR*_ the re-rupture rate. *De novo rupture rate*: Ever-ruptured nuclei (R+S) increased from 25% at day 11 post tamoxifen to 45% at day 14. Assuming first-order kinetics, the *de novo* rupture rate (*k*_*IR*_) was estimated from:

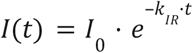, where *I*_0_ is initial intact fraction, which yielded *k*_*IR*_ ≈ 0.004 *h*^−1^.

#### Re-sealing rate

In live imaging, 23% of ruptured nuclei became resealed. Assuming first-order kinetics:

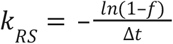, where *f* is the fraction resealed (23%), yielding *k*_*RS*_ ≈ 0.098 *h*^−1^.

#### Re-rupture rate

In live imaging, 40% of resealed nuclei became re-ruptured. Assuming first-order kinetics:

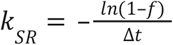, where *f* is the fraction resealed (40%), yielding *k*_*SR*_ ≈ 0.192 *h*^−1^.

Because resealing (0.098^−1^) and re-rupturing (0.192^−1^) occur much faster than *de novo* rupture (0.004h^−1^), the ruptured and resealed states rapidly approach a quasi-steady ratio 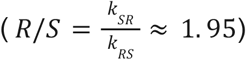, consistent with the observed proportions at day 14 post tamoxifen (1.74 = 33% ruptured over 19% resealed).

#### Estimation of rupture onset

Assuming that nuclei are initially all intact and that *de novo* ruptures begin at time *t*_0_, the cumulative fraction of nuclei that have ruptured is:

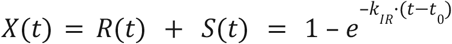

Solving this equation using the measured value with observed icGAS^+^ nuclei at day 14 (*X*(*t*) = 0. 52) yielded *t*_0_ ≈ 6. 7 days post tamoxifen.

The estimated parameters were used in the system of differential equations to simulate the temporal evolution of intact, ruptured, and resealed nuclei. The computation was performed using the *ode* function from the *deSolve* package in R.

### Immunofluorescence on mouse cardiac tissues

Hearts from mice expressing NLS-tdTomato and GFP-icGAS were fixed in 10% formalin, embedded in O.C.T. compound (Sakura Finetek, 4583), and cryo-sectioned at 12 µm thickness. Sections were blocked in a buffer containing 5% normal donkey serum, 1% non-fat milk, and 0.1% Triton X-100 in PBS, and then incubated overnight at 4°C with goat anti-desmin antibody (Invitrogen, PA5–19063P), followed by Alexa fluorophore-conjugated secondary antibodies for 1 hr at 37°C. Tissues were counterstained with DAPI, and then mounted with ProLong Glass Antifade Mountant (Invitrogen, P36984). Immunofluorescence signals were acquired using a Yokogawa CSU-W1 Sora spinning disk confocal microscope with 60x objectives, using channels for GFP, tdTomato, Alexa Fluor 647, and DAPI, corresponding to GFP-icGAS, NLS-tdTomato, desmin immunostaining and nuclei, respectively.

### Immunofluorescence on human cardiac tissues

Paraffin sections of fixed hearts were deparaffinized using xylene and ethanol and subjected to heat-induced antigen retrieval in a 10 mM citrate buffer (pH 6.0) for 5 min. Antigen-retrieved sections were blocked in a buffer containing 5% normal donkey serum, 1% non-fat milk, and 0.1% Triton X-100 in PBS, and then incubated overnight at 4°C with mouse anti-BANF1 antibody (Abnova, H00008815-M01) and goat anti-desmin antibody (Invitrogen, PA5–19063P), followed by Alexa fluorophore-conjugated secondary antibodies for 1 hr at 37°C. Cells were counterstained with DAPI, submerged in 0.25% Sudan Black B (Electron Microscopy Slides, #21610) in 70% Isopropanol for 10 min, and then mounted with ProLong Glass Antifade Mountant (Invitrogen, P36984). Immunofluorescence images of human myocardial sections were acquired using a Yokogawa CSU-W1 Sora spinning disk confocal microscope with 40x or 60x objectives (40x for normal hearts and 60x for ACTC1-variant heart and LMNA-R541C heart). To quantify BANF1 enrichment at nuclear tips, we first placed 50 seeds for ROIs at random positions in an image. For each seed, we selected the closest nucleus and created an ROI (see below). We manually drew a line as an ROI along the long axis of each nucleus, with endpoints positioned at the nuclear tips. The width of ROIs was set to 0.325 µm (2 pixels) for images acquired with 40x objectives or 0.321 µm (3 pixels) for images acquired with 60x objectives, so that a similar physical width was used across images. A BANF1 intensity profile along each ROI was extracted in Fiji using custom macros. For each nucleus, the first and last 10% of the intensity profile were defined as nuclear tips, and the central 80% of the line was defined as the nuclear body. For each segment, we computed mean BANF1 intensity. The stronger of the two tip measurements was used to compute nuclear-tip BANF1 enrichment (log_2_ tip/body) for each nucleus. Nuclei were classified as positive for BANF1 puncta when the enrichment ratio exceeded 2-fold (log_2_ tip/body ≥ 1).

### Immunoblot

Isolated cardiomyocytes were lysed in Urea Buffer (20 mM HEPES pH 7.4, 1 M NaCl, 8 M urea, protease and phosphatase inhibitors) for Lamin A/C and GAPDH immunoblotting. Proteins were extracted using pestle homogenization and sonication. Cell lysates were centrifuged at 15,000 g for 3 min at 4°C, and proteins in the supernatant were separated by SDS-PAGE. Proteins were transferred to a PVDF membrane. Membranes were blocked with nonfat milk. Primary antibodies used were rabbit anti-Lamin A/C antibody (Santa Cruz, sc-20681, 1:1000) and rabbit anti-GAPDH antibody (ABclonal, AC001, 1:5000). Secondary antibodies used were anti-rabbit IgG (H+L) DyLight 680 (Cell Signaling, #5366, 1:5000) and anti-rabbit IgG (H+L) DyLight 800 (Cell Signaling, #5151, 1:5000). Signals were detected in the Odyssey CLx Imager (LI-COR).

### RNA Polymerase II ChIP-seq

Preparation of fixed mouse cardiomyocytes for ChIP is described in the *Cardiomyocyte isolation* section. For human spike-in control, BJ-5ta fibroblasts (see *Cell culture*) were fixed with 1% paraformaldehyde for 15 min at room temperature. Detailed methods for chromatin preparation, immunoprecipitation, and library preparation are described in our previous paper ^54^. Fixed cardiomyocytes and BJ-5ta cells were sonicated in LB3Triton (100 mM NaCl, 1 mM EDTA, 0.5 mM EGTA, 1% Triton X-100, 0.1% sodium deoxycholate, 0.5% N-lauroyl sarcosine, 10 mM Tris-HCl, pH 8.0) using Bioruptor (high setting; 10 min for mouse cardiomyocytes and 5 min for human BJ-5ta cells). For each immunoprecipitation reaction, lysate corresponding to 2.6 µg of mouse DNA was mixed with 0.15 µg of human spike-in DNA in LB3Triton supplemented with protease inhibitor cocktail (Sigma-Aldrich, 539131) and phosphatase inhibitors (Abcam, ab201113). Ten percent of cleared lysate was saved as an input control. Lysates were incubated with rabbit anti-Pol II NTD antibody (Cell Signaling Technology, #14958) overnight at 4 °C. Immunocomplexes were captured by Dynabeads Protein G beads (Invitrogen, 10004D), washed, and eluted in Elution Buffer (1% SDS, 250 mM NaCl, 10 mM Tris-HCl pH 8.0, 1 mM EDTA). ChIP and input control samples were treated with RNase A, Proteinase K, and reverse-crosslinked. Purified DNA was used for sequencing library construction using NEBNext Ultra II DNA Library Prep Kit for Illumina (New England Biolabs, E7645L). Libraries were sequenced 100 bases on the paired-end mode on an Illumina NovaSeqX sequencer.

### ChIP-seq analysis

We used the mouse mm39 reference genome, the mouse Gencode vM27 basic gene annotation, the human GRCh39 reference genome, and the human Gencode v38 basic gene annotation for analysis. We used the “protein coding” gene annotation in Gencode vM27 annotation for gene-centric analyses (total 21,834 genes).

Pol II ChIP-seq reads (paired-end 100 bp) were processed by *fastp* ^55^ to remove duplicated reads and aligned to the mouse and human reference genomes individually using *Bowtie2* (ver 2.4.2) ^56^. Reads with mapping quality (MAPQ) greater than 20 were used in subsequent analyses. Normalization with spike-in control reads is described in the *Spike-in normalization* section below. Spike-in normalized Pol II ChIP-seq read coverage and depth-normalized input read coverage are used to visualize signals in the UCSC genome browser.

For metagene analysis, we generated 100 equally spaced bins in the gene body, 5 equally spaced bins in the 1-kb region immediately upstream of the transcription start site, and 10 equally spaced bins in the 2-kb region immediately downstream of the transcription end site. Only genes with the gene length greater than 1 kb and fully extendable to the upstream and downstream regions were used in the subsequent analysis (21,177 genes). For each bin, a mean of spike-in normalized per-base coverage was computed for each Pol II ChIP-seq replicate using *Bedtools* ^*57*^, and then a mean across the genes was used in visualization.

For gene level analyses, we computed Pol II ChIP coverage for each gene body. Gene-body Pol II ChIP coverage was the sum of spike-in normalized per-base read coverage within a gene body, normalized by the gene length in kb (**Table S1**). We removed genes with the size smaller than or equal to 1 kb as well as those with a minimum Pol II coverage below 100 across all replicates, which exhibited essentially no discernable Pol II ChIP-seq signals (**Fig. S5B**). This resulted in 11,942 Pol II-bound protein-coding genes. Log_10_-transformed Pol II coverage in the 11,942 genes for 3 biological replicates for each genotype was used in principal component analysis and differential enrichment analysis using *limma* ^58^. Genes with adjusted P-value < 0.05 were defined as differentially enriched (4 Pol II-gained and 1,759 Pol II-lost genes; **Table S1**). Enrichment of Gene Ontology terms within Biological Process, Molecular Function, and Cellular Component classes was computed using *Metascape* ^59^. For comparison with gene expression, we used whole heart RNA-seq data (GEO accession ID: GSE241577) and heart single-nucleus RNA-seq data (GEO accession ID: GSE241587) from wild-type and *Lmna*^*CKO*^ mice ^7^.

### Spike-in normalization

Human spike-in chromatin was used to normalize differences in ChIP efficiency across experiments. For each ChIP-seq experiment *i*, we computed a scaling factor based on spike-in reads (see below) and applied it linearly to per-base mouse ChIP-seq read coverage by multiplication. Assuming that the human spike-in chromatin is added in strict proportion to mouse chromatin, the scaling factor would be inversely proportional to the number of human-aligned ChIP-seq reads, *hChIP*_*i*_. In practice, however, the spike-in fraction can vary between reactions due to pipetting and chromatin quantification errors, which directly affect *hChIP*_*i*_. To account for this, we estimated the spike-in fraction *f* in each reaction using the corresponding input control: *f*_*i*_ = *hINPUT*_*i*_ /(*hINPUT*_*i*_ + *mINPUT*_*i*_), where *hINPUT*_*i*_ and *mINPUT*_*i*_ are the total input reads aligned to the human and mouse genome, respectively. The scaling factor (*SF*) for each ChIP-seq reaction was then defined as: 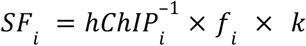, where *k* is a constant (set to 10^6^ for convenience).

Mathematically, let the mouse DNA-bound Pol II (“epitope”) concentration in ChIP reaction *i* be *M*_*i*_ and the human spike-in epitope concentration be *H*_*i*_. Let ChIP efficiency be *E*_*i*_. Let the observed total mouse and human ChIP read counts be *mChIP*_*i*_ and *hChIP*_*i*_, respectively, and the observed total mouse and human input read counts be *mINPUT*_*I*_ and *hINPUT*_*i*_, respectively. ChIP read counts are proportional to the product of epitope concentration and ChIP efficiency:

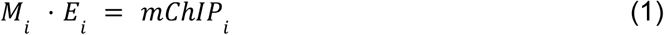

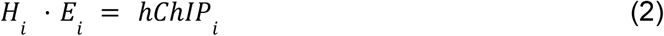

Combining (1) and (2):

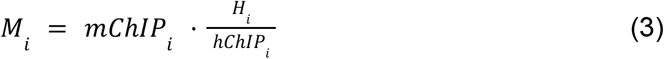

The human spike-in epitope concentration *H*_*i*_ depends on the spike-in proportion, which is estimated from input reads. Let *k* denote the desired spike-in level across experiments:

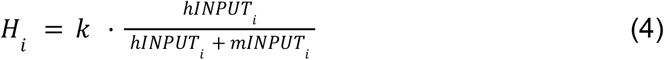

Substituting (4) into (3):

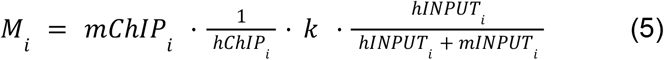

At a mouse genomic location *j*, (5) gives:

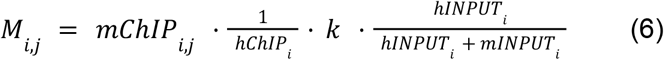

(6) can be rewritten as:

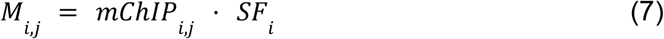

where scaling factor (SF) is:

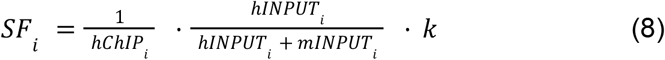

Read counts and scale factors are listed in **Fig. S5A**. Spike in-normalized ChIP-seq read coverage data are available in GEO under accession ID **GSE325163**. Applying a scaling factor to per-base coverage was performed using *deepTools* ^60^.

### Statistical analysis

All statistical analyses were performed in R (version 4.4.2). Statistical analyses were designed to account for the hierarchical structure of the data, in which multiple nuclei were measured from one slide which corresponded to one biological replicate of a mouse. Because nuclei from the same mouse are not independent and share common biological factors (e.g. extent of Lamin A/C depletion and AAV transduction efficiency) and technical factors (e.g., staining efficiency and tissue condition), we performed statistical analyses that accounted for this non-independence by treating mouse (slide) as the unit of clustering. Depending on the type of input and analysis, we used either generalized linear mixed-effects models (GLMMs) or linear regression models with mouse-clustered robust standard errors. GLMMs were used for binary or proportion data (e.g., nuclear states or LEMD2-associated nuclei), whereas linear regression with cluster-robust standard errors was used for continuous measurements (e.g., fluorescence intensities). These approaches allowed appropriate estimation of variability at the level of mice rather than individual nuclei and avoided underestimation of biological variability that could lead to false-positive results. In analyses performed at the sample level (i.e., one value per sample or image), where clustering of nuclei was not present, conventional statistical tests were used. Unless otherwise stated, statistical tests were two-sided. Data are presented as mean ± standard error (SE) unless otherwise indicated. Below, we describe details of statistical analyses:

#### Comparison of ruptured versus resealed nucleus proportions

To compare the proportions of ruptured and resealed nuclei in *Lmna*^*CKO*^ mice, we used the numbers of ruptured nuclei in each sample to retain information about the total number of nuclei analyzed in each sample. We used the data set that described the nuclear state in binary (ruptured or resealed) for each nucleus for each sample. A binomial GLMM with a logit link was fitted. In this model, the responses were the proportion of ruptured nuclei, and the random effect was the mouse identity, which accounted for differences between mice and the non-independence of nuclei from the same sample. The model estimated the mean probability for the ruptured nuclear state and its variance. Statistical significance was evaluated by testing against the null hypothesis that ruptured and resealed nuclei occur at equal frequency (i.e., probability of rupture = 0.5). The null distribution was modeled using parametric bootstrap (2,000 simulations). We used the R package *lme4* for this analysis.

#### Comparison of gamma-H2AX, BrU, and Pol II intensities between different nuclear states

To compare gamma-H2AX, BrU, and Pol II intensities across nuclear states, we used normalized fluorescence intensities for each nucleus (see above). Linear regression models were fitted with nuclear states as categorical predictors and normalized fluorescence intensities as the response. Mouse-level clustering was accounted for using cluster-robust standard errors. The model estimated mean intensity values (estimated marginal means) for each nuclear state and their cluster-adjusted standard errors. In this context, “cluster adjustment” means that nuclei from the same mouse were treated as a group. Pairwise differences between nuclear states were evaluated using *t*-tests based on these estimates, and corresponding p-values were calculated for the null hypothesis of no difference. Multiple comparisons were adjusted using the Tukey method. We used R packages *sandwich, lmtest, emmeans*, and *stats* for this analysis.

#### Relationship between gamma-H2AX, BrU, or Pol II intensities and nuclear tdTomato

To analyze the relationship between tdTomato intensity and nuclear signal intensities (γ-H2AX, BrU, and Pol II), we used normalized fluorescence intensities for each nucleus as inputs (see above). Linear regression models were fitted with normalized tdTomato intensity as a continuous predictor and normalized nuclear intensities as a response. Statistical significance of the association was assessed using *t*-tests for the regression coefficient, with cluster-robust standard errors clustered by mouse identity. We used R packages *sandwich, clubSandwich*, and *Imtest* for this analysis.

#### LEMD2- and VPS4-associated nucleus proportion between ruptured and resealed nuclei

To compare the proportions of LEMD2- or VPS4-associated nuclei between ruptured and resealed nuclei, we used the data set that described the LEMD2- or VPS4-association state in binary (associated or not) and the nuclear state in binary (ruptured or resealed) for each nucleus for each sample. Separate analyses were conducted for LEMD2 and VPS4. We used a binomial GLMM with a logit link, with the nuclear state as a categorical predictor, the association state as a response, and mouse identity as a random effect. The model estimated the probability of LEMD2- or VPS4-associated nuclei and the variance. Statistical significance was assessed by Wald z-tests. The null hypothesis was that there is no difference in the proportion of associated nuclei between nuclear states. We used the R package *lme4* for this analysis.

#### Relationship between LEMD2- or VPS4-association and nuclear tdTomato

For visualization of the relationship between tdTomato intensity and LEMD2- or VPS4-association, a generalized additive model (GAM) with a smoothing spline was used to fit the data. The smoothing parameter (k = 5) was chosen to limit overfitting. Shaded areas represent 95% confidence intervals. This analysis was used for descriptive purposes only and not for statistical inference. To analyze the relationship between tdTomato intensity and LEMD2- or VPS4-association, we used a data set that described the LEMD2- or VPS4-association state in binary (associated or not) and normalized *tdTomato* intensities for each nucleus for each sample (see above). Binomial generalized GLMMs with a logit link were fitted to model the probability of a nucleus being protein-positive, with normalized tdTomato intensity as a continuous predictor and mouse identity as a random effect. Statistical significance of the association was evaluated using Wald z-tests for the tdTomato coefficient in the model. We used the R package *lme4* for this analysis.

#### Comparison of ruptured and resealed nucleus composition between genotypes

To compare the relative frequency of ruptured and repaired nuclei between *Lmna*^*CKO*^ and *Lmna*^*CKO*^*;VPS4*^*EQ*^, we modeled nuclear state (ruptured vs repaired) as a binary outcome at the level of individual nuclei. Binomial GLMMs with a logit link were fitted with genotypes as a categorical predictor, the nuclear state binary outcome as a response, and mouse identity as a random effect. Statistical significance was assessed using Wald z-tests based on the effect of genotype in the model. A one-sided test was used to evaluate whether the VPS4-EQ increased the probability of nuclei being ruptured relative to being resealed compared to the control genotype. The R package *lme4* was used for this analysis.

#### Comparison of percentages of cGAS^+^ nuclei between genotypes

To compare the percentages of cGAS^+^ nuclei between *Lmna*^*CKO*^ and *Lmna*^*CKO*^*;VPS4*^*EQ*^, we used the percentages of cGAS^+^ nuclei in each sample. Genotype differences were assessed by the unpaired Welch’s t-test to test the null hypothesis of no difference. We used the R package *stats* for this analysis.

#### Comparison of fibrotic area between genotypes

Trichrome area fraction was analyzed as a proportion and logit-transformed to place values on an unbounded scale suitable for parametric analysis. To avoid undefined values at 0 or 1, proportions were bounded slightly away from these limits (minimum 10^−6^, maximum 1–10^−6^). Genotype differences were assessed by one-way ANOVA with Tukey’s post hoc correction. The R package *stats* was used for this analysis.

#### Comparison of BANF1-positive nucleus fraction between human cardiac specimens

We used the percentages of BANF1-positive nuclei in each image. Statistical differences between human cardiac specimens were evaluated based on the Kruskal-Wallis test, followed by pairwise Wilcoxon rank-sum tests with Holm adjustment for multiple comparisons. The R package *stats* was used for this analysis.

#### Gene expression change of Pol II-lost genes versus other genes

We used RNA-seq log_2_ fold change (*Lmna*^*CKO*^/wild type) for Pol II-lost genes and all other genes as inputs for two-sample Kolmogorov-Smirnov test. RNA-seq data were either from bulk whole-heart RNA-seq or pseudo-bulk cardiomyocyte RNA-seq. The null hypothesis was that the two gene types came from the same log_2_ fold change distribution. The R function *ks*.*test* was used.

---

**Figure S1.**
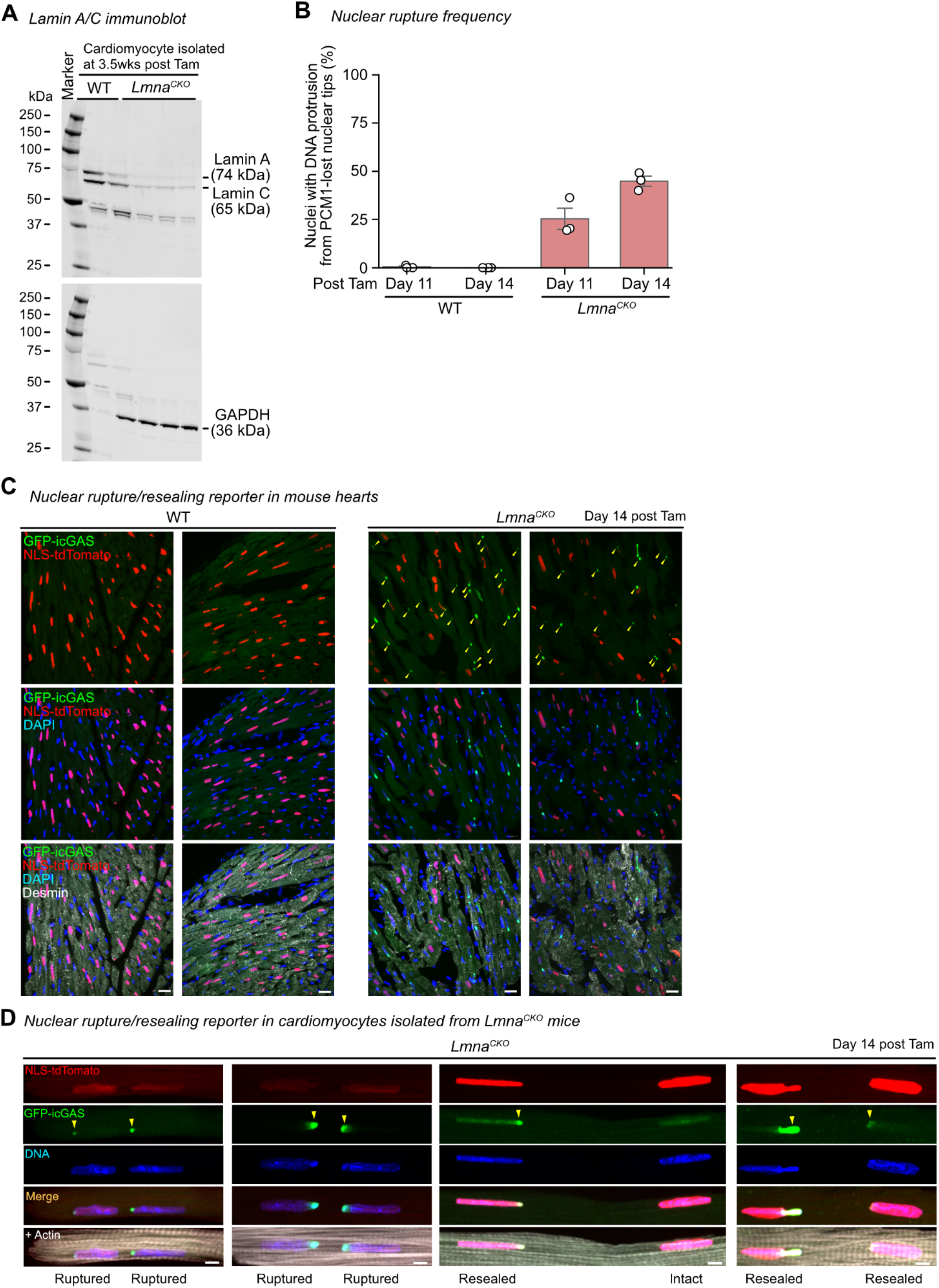
Identification of intact, ruptured and resealed nuclei in *Lmna*^*CKO*^ hearts. **A)** Immunoblot for Lamin A/C (top) and GAPDH (bottom) in cardiomyocytes isolated from wild-type (WT) or *Lmna*^*CKO*^ hearts at 3.5 weeks post tamoxifen. **B)** Percentage of nuclei with DNA protrusion from PCM1-lost nuclear tip at day 11 and 14 post tamoxifen. Circles: percentages in 3 biological replicates. Column: mean of the replicates. Error bar: standard error. Nuclear count: 199 (WT, day 11), 176 (WT, day 14), 381 (*Lmna*^*CKO*^, day 11), and 351 (*Lmna*^*CKO*^, day 14) from 3 mice per genotype for each time point. **C)** Additional images of heart sections of mice expressing NLS-tdTomato and GFP-icGAS, co-stained for Desmin and DNA (DAPI). Day 14 post tamoxifen. Arrowheads: icGAS puncta at rupture sites. Scale bar: 5 μm. **D)** Additional images of cardiomyocytes isolated from mice expressing NLS-tdTomato and GFP-icGAS at day 14 post tamoxifen. Arrowheads: GFP-icGAS puncta at rupture sites. Scale bar: 5 μm.

**Figure S2.**
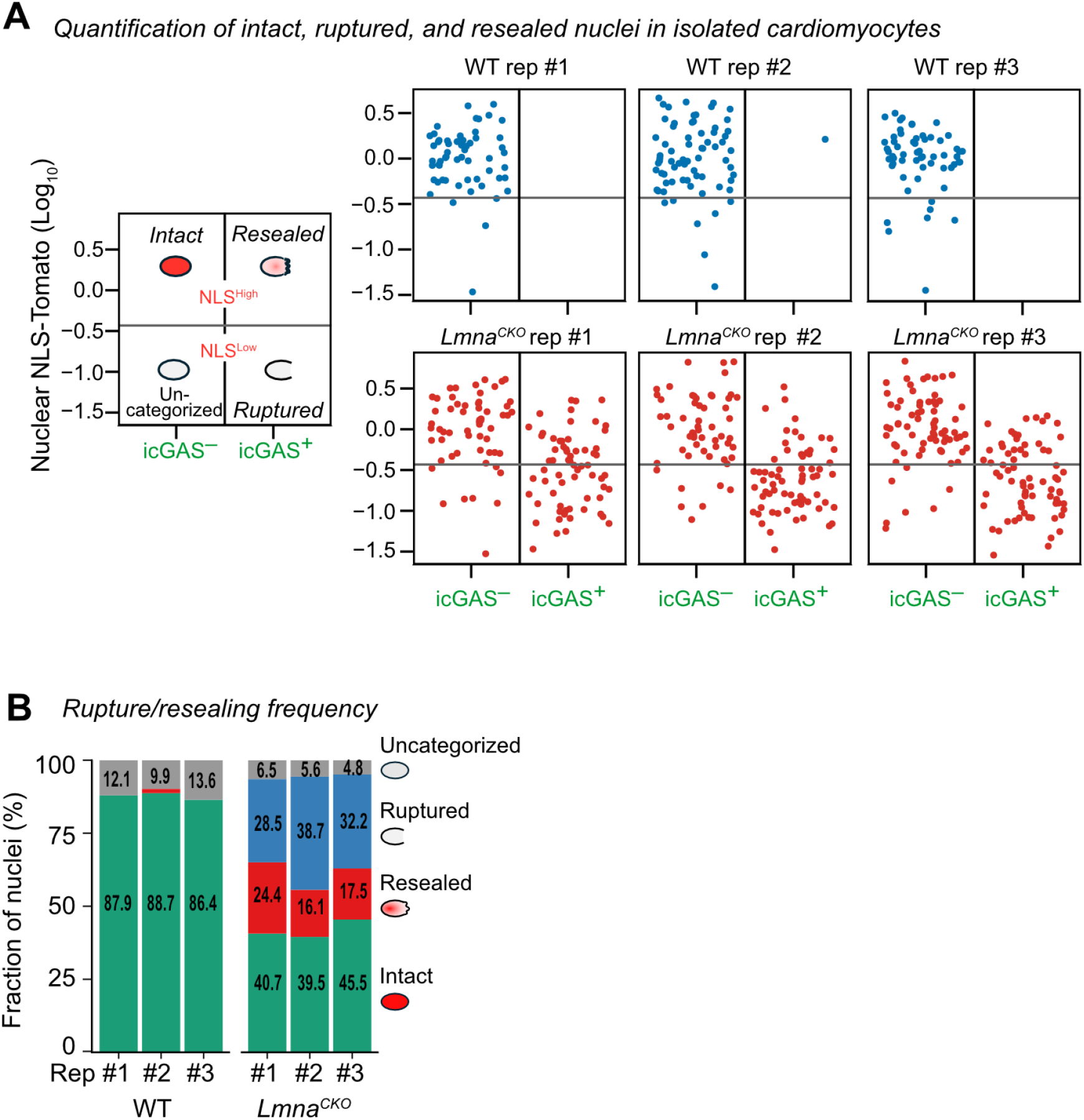
Quantification of intact, ruptured and resealed nuclei. **A)** Classification of cardiomyocyte nuclei by the presence or absence of GFP-icGAS punctum at nuclear tips (x-axis) and nuclear NLS-tdTomato intensity (y-axis) in individual mice. Horizontal line: mean minus one standard deviation of nuclear tdTomato intensity in icGAS^−^ nuclei from all replicates. **B)** Proportion of intact, ruptured, resealed, and uncategorized nuclei in individual mice. N=3 mice per genotype.

**Figure S3.**
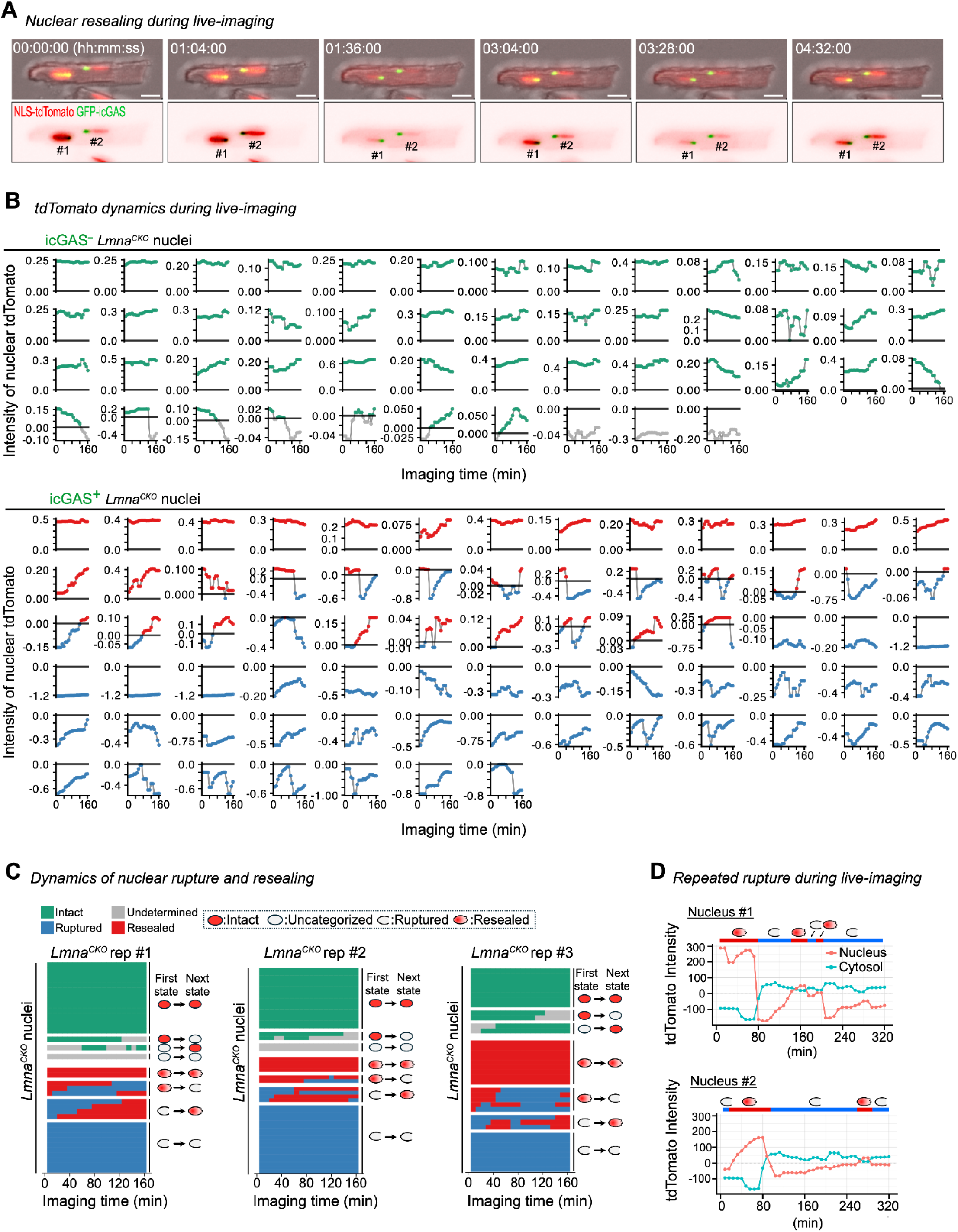
Time-lapse imaging of nuclear rupture and resealing. **A)** A representative *Lmna*^*CKO*^ cardiomyocyte undergoing nuclear re-rupture and resealing. Fluorescent signals are shown with (top) or without (bottom) bright-filed image. Images at 00:00:00, 01:36:00, 03:04:00, and 03:28:00 are also shown in main Figure 1G. Scale bar: 20 μm. **B)** Nuclear tdTomato intensity profiles during 2.5-hr time-lapse imaging for all 121 *Lmna*^*CKO*^ cardiomyocyte nuclei used in this study. Top: icGAS^−^ nuclei (i.e. intact and uncategorized nuclei). Bottom: icGAS^+^ nuclei (i.e. ruptured and resealed nuclei). Lines are colored by nucleus state at the indicated time (green, intact; blue, ruptured; red, resealed; gray, uncategorized). Data are centered to the mean minus standard deviation of tdTomato intensity in icGAS^−^ nuclei at the indicated time from the same mouse. **C)** Nuclear state dynamics during 2.5-hr time-lapse imaging in three biological replicates (*Lmna*^*CKO*^ mice). Row: individual nuclei (replicate 1, 36; replicate 2, 48; replicate 3, 37). Column: imaging time. **D)** Dynamics of nuclear and cytoplasmic tdTomato intensities in two representative nuclei in one *Lmna*^*CKO*^ cardiomyocyte undergoing repetitive nuclear re-rupture and resealing. The nuclear intensity is scaled 1/10 so that both nuclear and cytoplasmic intensities are in the y-axis range. The images of this cardiomyocyte are shown in **(A)**.

**Figure S4.**
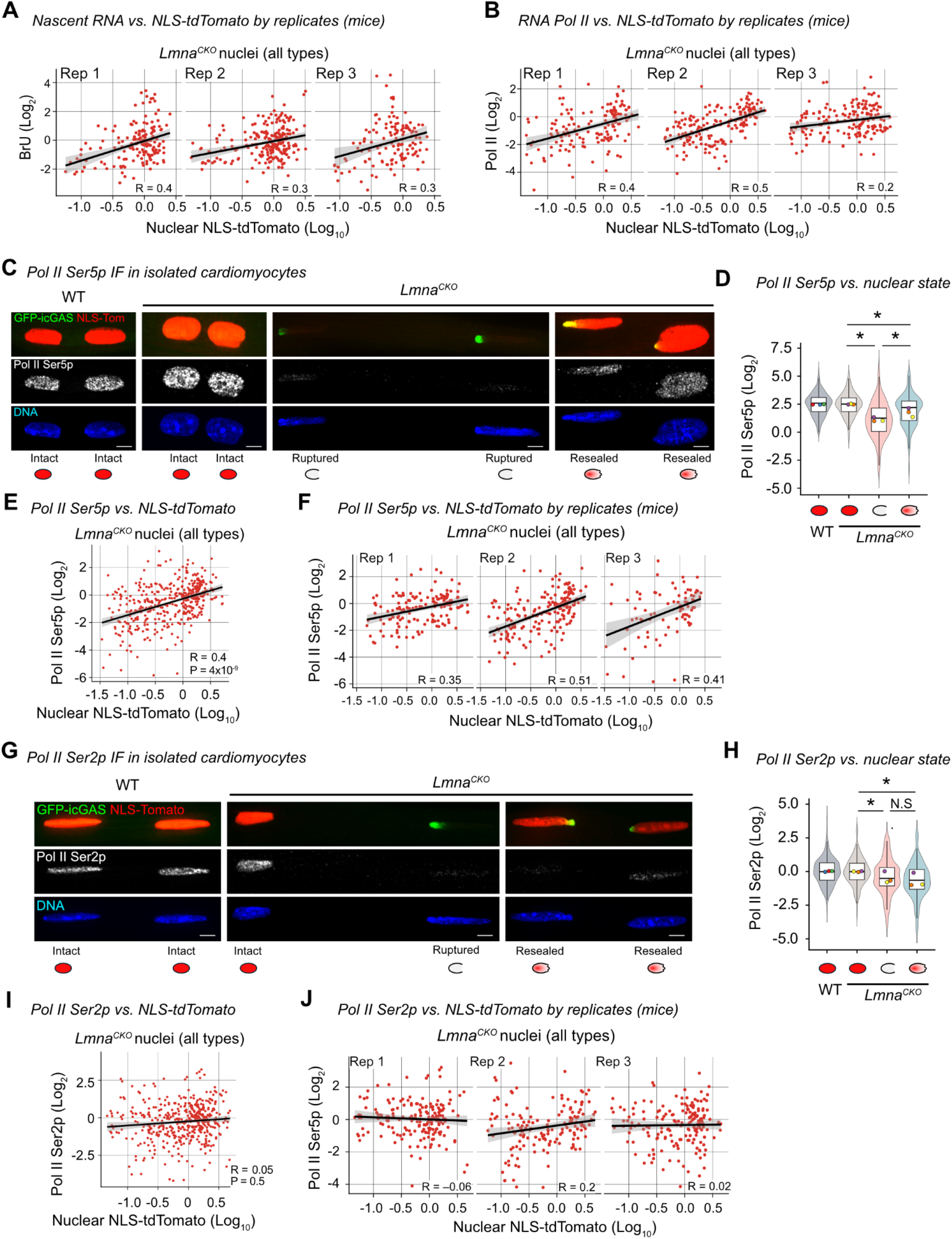
Quantification of transcription and RNA Pol II in cardiomyocyte nuclei. **A)** Relationship between nuclear BrU intensity and nuclear NLS–tdTomato intensity in cardiomyocytes for each of 3 *Lmna*^*CKO*^ mice. Line: simple linear regression fit with 95% confidence interval. R: Pearson’s correlation coefficient. All data in **Figure S4** are derived from mice at day 14 post tamoxifen. **B)** Relationship between nuclear RNA Pol II intensity and nuclear NLS-tdTomato intensity in cardiomyocytes for each of 3 *Lmna*^*CKO*^ mice. Other details as in **(A)**. **C)** Immunofluorescence for Pol II CTD phospho-Ser5 (Pol II Ser5p) in isolated cardiomyocytes expressing NLS-tdTomato and GFP-icGAS. Scale bar: 5 μm. **D)** Nuclear Pol II Ser5p intensity by nuclear states. Density and box plots (interquartile range): signal distribution of all nuclei. Circles: mean intensity within individual biological replicates (color coded). Asterisks: P < 0.05 from t-tests on linear regression-estimated means with mouse-clustered standard errors. Underlying data: 218 intact nuclei from 3 WT mice, 187 intact, 111 ruptured, 91 resealed nuclei from 3 *Lmna*^*CKO*^ mice. **E, F)** Relationship between Pol II Ser5p intensity and NLS–tdTomato intensity in nuclei of *Lmna*^*CKO*^ cardiomyocytes. **(E)** 424 nuclei from three biological replicates (mice). **(F)** Analysis within individual biological replicates. Other details as in **(A)**. **G)** Immunofluorescence for Pol II CTD phospho-Ser2 (Pol II Ser2p) in isolated cardiomyocytes expressing NLS-tdTomato and GFP-icGAS. Scale bar: 5 μm. **H)** Nuclear Pol II Ser2p intensity by nuclear states. Underlying data: 198 intact nuclei from 3 WT mice, 233 intact, 101 ruptured, 131 resealed nuclei from 3 *Lmna*^*CKO*^ mice. N.S.: not significant. Other details as in **(D)**. **I, J)** Relationship between Pol II Ser2p intensity and NLS-tdTomato intensity in nuclei of *Lmna*^*CKO*^ cardiomyocytes. **(I)** 517 nuclei from three biological replicates (mice). **(J)** Analysis within individual biological replicates. Other details as in **(A)**.

**Figure S5.**
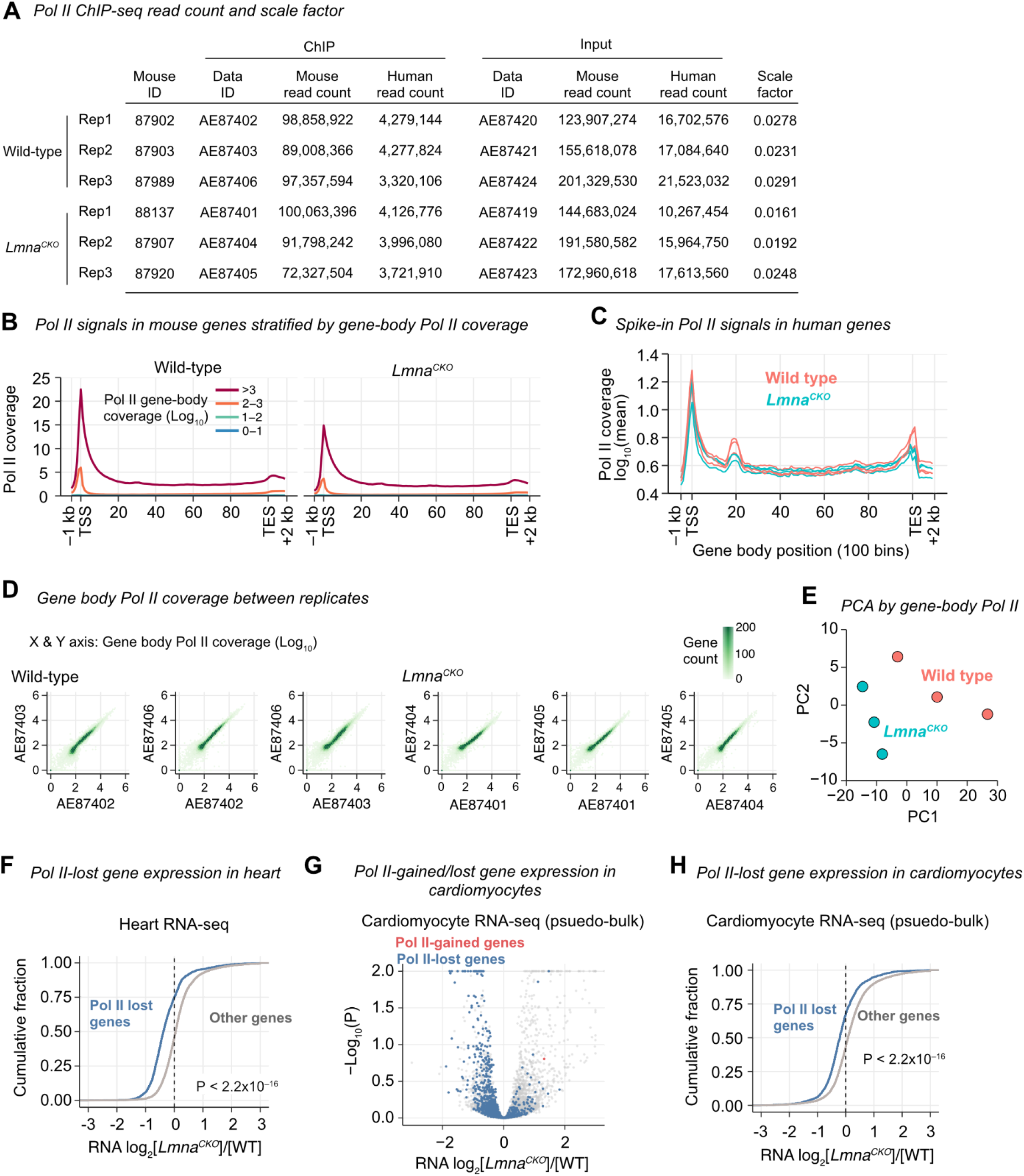
RNA Pol II ChIP-seq in isolated cardiomyocytes. **A)** Number of Pol II ChIP-seq and input sequencing reads aligned to the mouse genome (experimental) or the human genome (spike-in control). Scale factors are computed by spike-in control reads and sequencing depths and used to normalize Pol II ChIP-seq signals. All data in **Figure S5** are derived from mice at 2 weeks post tamoxifen. **B)** Pol II ChIP-seq read coverage in all mouse genes stratified by gene-body Pol II coverage. Genes with Pol II coverage greater than or equal to 100 (2 in Log_10_) were considered Pol II-bound (11,942 genes). **C)** Pol II ChIP-seq read coverage in all human genes, derived from the spike-in control chromatin. **D)** Gene-body Pol II ChIP-seq read coverage between every pair of biological replicates. **E)** Principal Component Analysis (PCA) of gene-body Pol II coverage in 11,942 Pol II-bound protein-coding genes. **F)** Cumulative fraction of 1,759 Pol II-lost genes and all other genes (y-axis) along the scale of differential gene expression between *Lmna*^*CKO*^ hearts and wild-type hearts (x-axis). P, Kolmogorov-Smirnov test *p*-value comparing log_2_ fold change of gene expression between Pol II-lost genes and all other genes. **G)** Gene expression state of Pol II-lost genes, Pol II-gained genes, and all other genes in the cardiomyocyte population in *Lmna*^*CKO*^ (n=3) versus WT (n=3) hearts derived from single-nucleus RNA-seq in En et al. 2024. P, DESeq2 *p*-value. **H)** Same as F, but along the scale of differential gene expression between *Lmna*^*CKO*^ and wild-type pseudo-bulk cardiomyocytes from the single-nucleus RNA-seq. P, Kolmogorov-Smirnov test *p*-value comparing log_2_ fold change of gene expression between Pol II-lost genes and all other genes.

**Figure S6.**
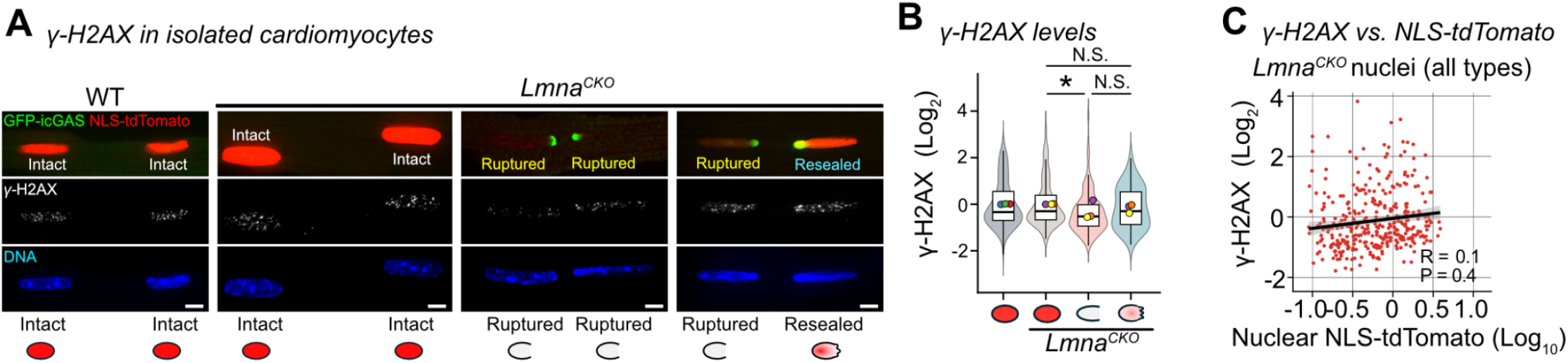
DNA damage analysis in *Lmna*^*CKO*^ cardiomyocyte nuclei. **A)** Immunofluorescence for gamma-H2AX in cardiomyocytes expressing NLS-tdTomato and GFP-icGAS isolated from mice at day 14 post tamoxifen. Texts and icons below images indicate nuclear states. Scale bar: 5 μm. **B)** Nuclear gamma-H2AX intensity by nuclear states. Density and box plots (interquartile range): signal distribution of all affiliated nuclei. Circles: mean intensity within individual biological replicates (color coded). Statistics: P < 0.05 (*) or P≥0.05 (N.S.) from t-tests on linear regression-estimated means with mouse-clustered standard errors. Underlying data: 164 intact nuclei from 3 WT mice, 152 intact, 116 ruptured, 50 resealed nuclei from 3 *Lmna*^*CKO*^ mice. **C)** Relationship between gamma-H2AX intensity and NLS-tdTomato intensity in nuclei of *Lmna*^*CKO*^ cardiomyocytes (340 nuclei from 3 mice). Line: simple linear regression fit with 95% confidence interval. R: Pearson’s correlation coefficient. P: t-test *p*-value on linear regression-estimated means with mouse-clustered standard errors.

**Figure S7.**
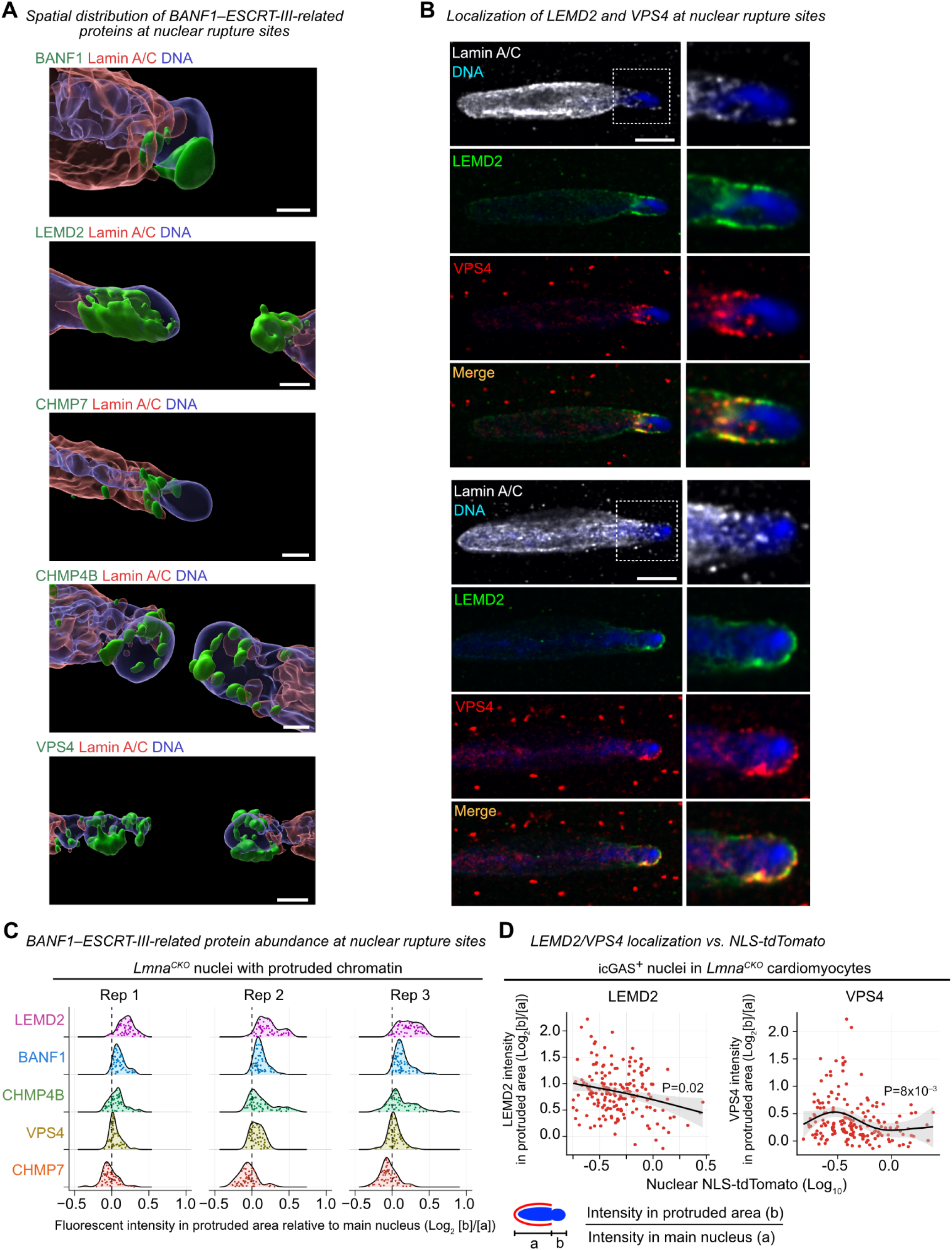
Localization of BANF1–ESCRT-III proteins at nuclear rupture sites. **A)** Localization of BANF1–ESCRT-III pathway proteins on protruded chromatin at nuclear rupture sites, visualized by three-dimensional image segmentation of raw image signals. Scale bar: 5 μm. All data in **Figure S7** are derived from mice at day 14 post tamoxifen. **B)** Localization of LEMD2 and VPS4 on protruded chromatin in high magnification. Left, two representative nuclei in *Lmna*^*CKO*^ cardiomyocytes. Right, magnified images of areas indicated by boxes in left images. Scale bar: 5 μm. **C)** Immunofluorescent intensity of BANF1–ESCRT-III pathway proteins at protruded chromatin area (“b” in schematic) relative to main nuclear area (“a” in schematic) in *Lmna*^*CKO*^ cardiomyocytes. Data points: log_2_[b]/[a] ratio of individual protruded chromatin. Line: density estimate of the data points. **D)** Relationship between relative LEMD2 or VPS4 intensity in protruded chromatin area (y-axis, Log_2_[b]/[a] ratio) and nuclear tdTomato intensity (x-axis) among all icGAS^+^ nuclei (i.e. ruptured and resealed nuclei) in *Lmna*^*CKO*^ cardiomyocytes. Line: non-linear fit with 95% confidence interval. P: Wald z-test *p*-value on binomial GLMMs estimating the probability of association. Nucleus count: LEMD2 n=170, VPS4 n=177 pooled from three mice.

**Figure S8.**
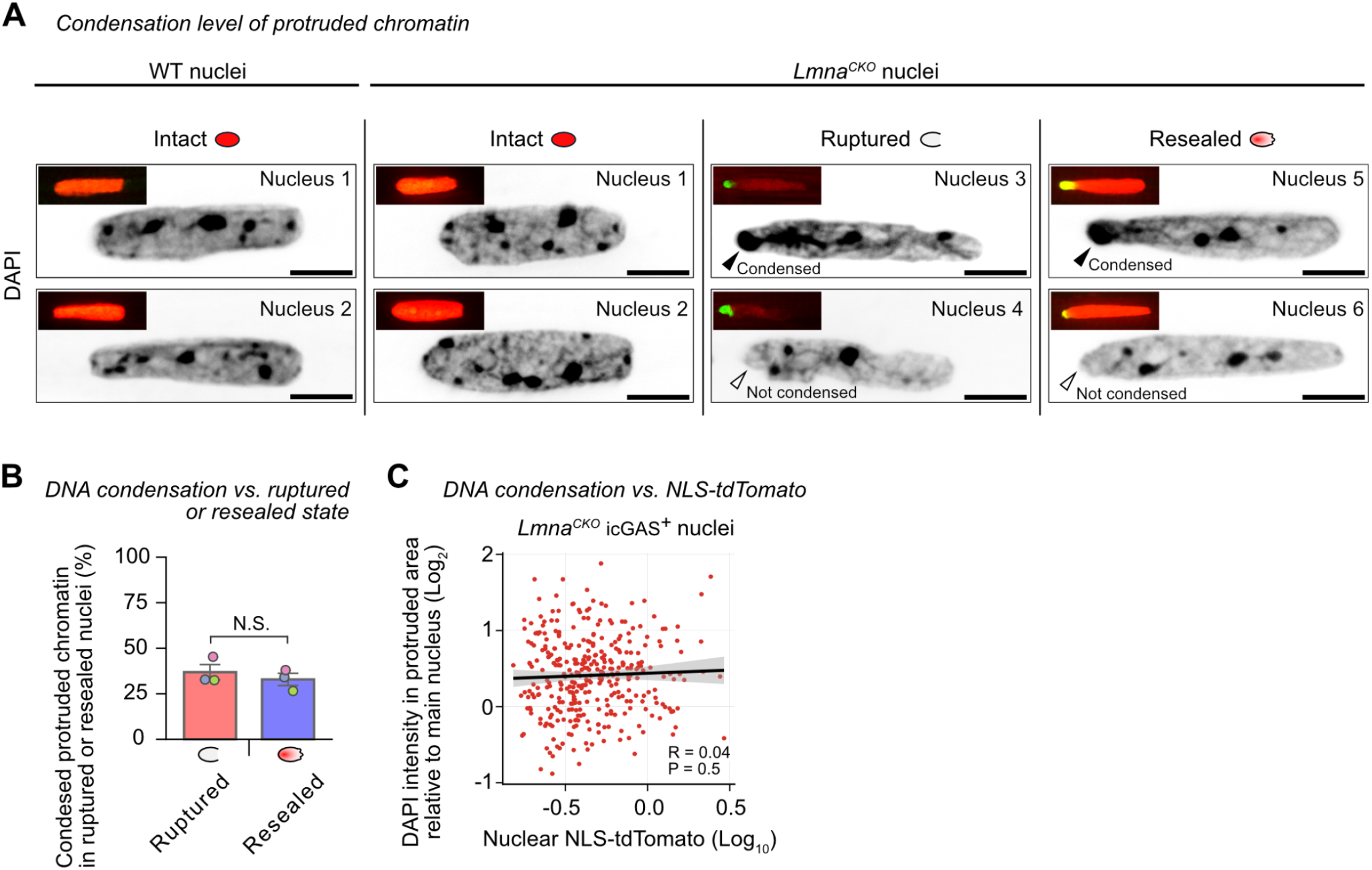
Analysis of DNA condensation of protruded chromatin. **A)** DAPI signals in intact, ruptured, and resealed nuclei in *Lmna*^*CKO*^ cardiomyocytes. For ruptured and resealed nuclei, condensed (top) and uncondensed protruded chromatin (bottom) are shown. Inset: the same nucleus with GFP-icGAS and NLS-tdTomato reporters. All **Figure S8** data are from cardiomyocytes isolated from mice and immediately fixed for experiments at day 14 post tamoxifen. **B)** Fraction of *Lmna*^*CKO*^ ruptured or resealed nuclei with condensed protruded chromatin. Statistics: Wald z-tests in binomial generalized linear mixed model (N.S.: p-value ≥0.05). **C)** Relationship between DAPI intensity in protruded chromatin area (y-axis) and nuclear tdTomato intensity (x-axis) for all icGAS^+^ nuclei (i.e. ruptured and resealed nuclei) in *Lmna*^*CKO*^ cardiomyocytes. R: Pearson’s correlation coefficient. P: t-test *p*-value on linear regression-estimated means with mouse-clustered standard errors. Nucleus count: n=348 (205 ruptured, 143 resealed) pooled from three mice.

**Figure S9.**
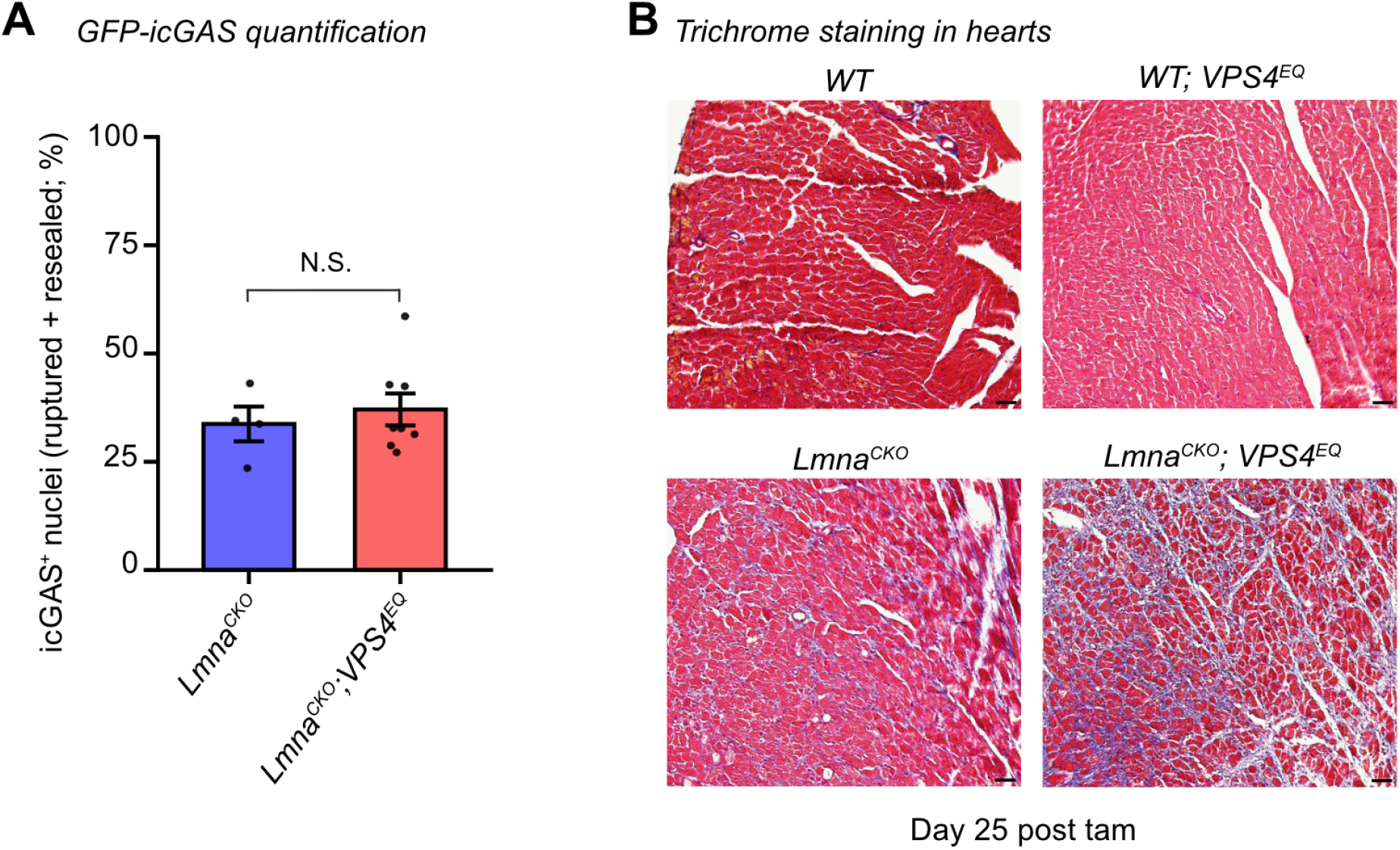
Dominant-negative VPS4-EQ expression in *Lmna*^*CKO*^ mice. **A)** Fraction of icGAS^+^ nuclei (i.e. ruptured and resealed nuclei) among all nuclei in cardiomyocytes isolated from *Lmna*^*CKO*^ (n=4) or *Lmna*^*CKO*^*;VPS4*^*EQ*^ (n=8) mice at day 14 post tamoxifen. Bar: mean +/– standard error. P: unpaired one-tailed Welch’s t-test. **B)** Masson’s trichrome staining of heart sections at day 25 post tamoxifen. Scale bar: 20 μm.

**Figure S10.**
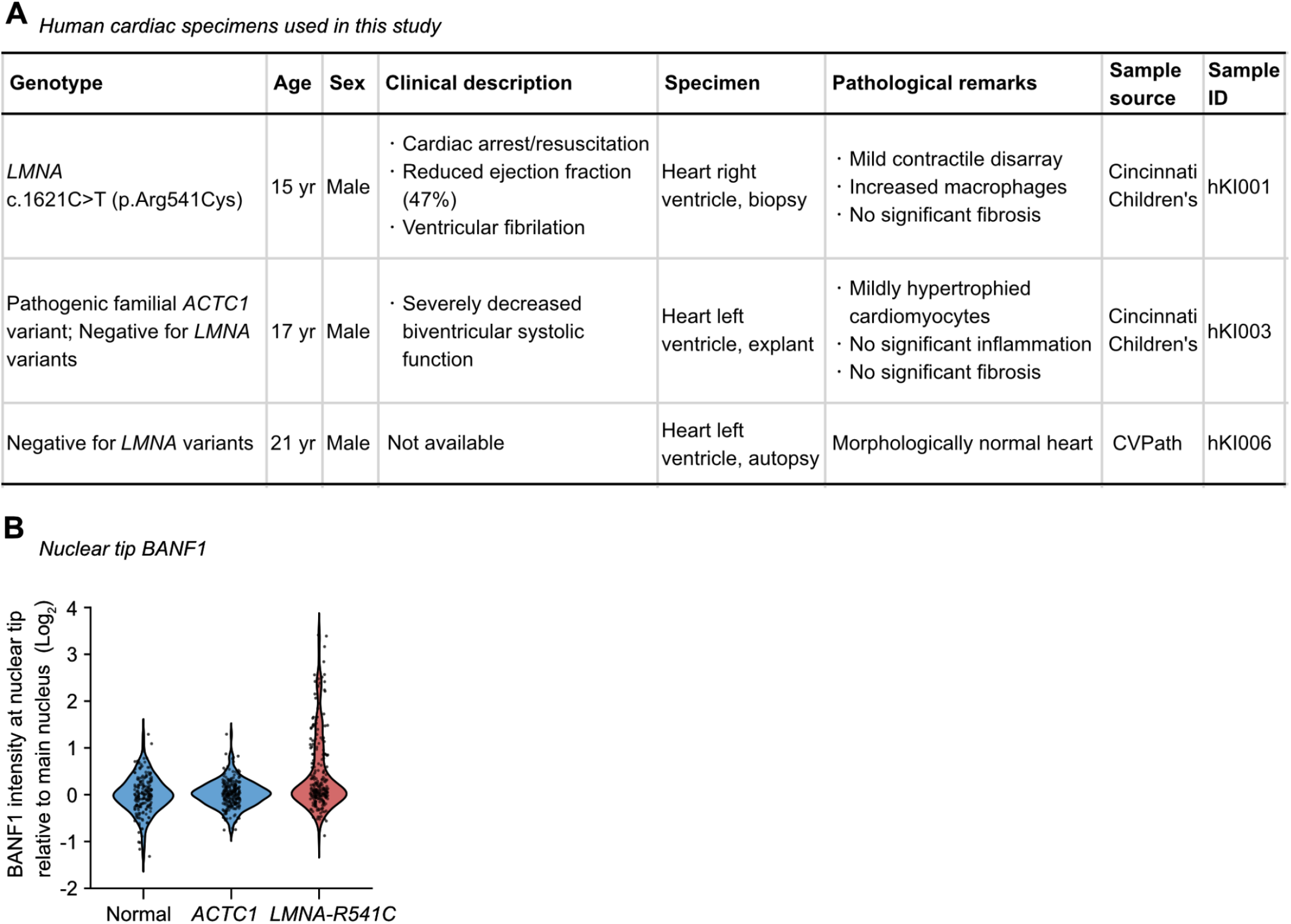
Human cardiac specimens used in this study. **A)** Table lists specimens used in BANF1 immunofluorescence in Figure 4. **B)** BANF1 intensity at the tip of individual nuclei in cardiomyocytes. Nuclear count: human normal heart n=200 from 4 specimen fields, human *ACTC1*-mutant heart n=250 from 5 specimen fields, and *LMNA* p.R541C heart n=250 from 5 specimen fields.

